# Essential Roles of RodA Peptidoglycan Polymerase and PBP2 Transpeptidase in Expression of Cell Wall-Spanning Supramolecular Organelles and Modulating *Salmonella* Virulence

**DOI:** 10.1101/2024.07.01.601524

**Authors:** Anne C. Doble, Bethany C Gollan, John Clark-Corrigall, David M. Bulmer, Richard A Daniel, Pietro Mastroeni, C. M. Anjam Khan

## Abstract

The increased spread of multidrug-resistant bacteria no longer sensitive to commonly used antibiotics poses a major threat to human health. The search for potential new drug targets is critical in disease control and prevention. Whilst several components of the cell wall synthesis machinery are already targeted by beta-lactam antibiotics, other elements of this machinery present opportunities for novel drug targets. Landmark studies revealed RodA exhibits peptidoglycan polymerase activity in *Bacillus subtilis* and *Escherichia coli,* highlighting RodA as a prime for the next generation of antimicrobial drugs. However, the role of RodA in virulence remains unexplored. Through targeted mutagenesis, virulence gene reporter assays, and phenotypic screening, we demonstrate that the presence of RodA or PBP2, is intrinsically linked to the regulation of virulence gene expression in *Salmonella*. Specifically, deletion of either of these components causes both disruption in cell morphology and a complete downregulation in major cell invasion-associated virulence factors *in vitro,* and attenuated virulence *in vivo*. Significantly, this study highlights the importance of RodA and PBP2 in both the biology and virulence of an important bacterial pathogen, identifying them as promising targets for developing new antibiotics.

## Introduction

*Salmonella enterica* cause a broad spectrum of disease in humans ranging from mild gastroenteritis to life-threatening typhoid fever, paratyphoid fever and invasive non-typhoidal salmonellosis (iNTS) (1–3). *Salmonella* Typhi causes up to an estimated 9M cases of typhoid fever per year and 110,000 deaths annually worldwide(4). Following apparent resolution, relapses occur in up to 10% of patients (5, 6) and chronic carriage in up to 6% of treated individuals(7). Invasive-Non Typhoidal Salmonellosis (iNTS) leads to sepsis, which can be fatal and cause relapsing infections in children, immune-compromised adults (e.g. individuals with HIV infection) and patients co-infected with malaria(3, 8). Invasive non-typhoidal salmonellosis (iNTS) cause an estimated 3.4M illnesses with 680,000 deaths annually and very high case-fatality ratios (20%). These infections are also difficult to clear with antibiotics. There are currently no licensed vaccines for paratyphoid fever and iNTS strains of *Salmonella*(3, 9). The increasing spread of multidrug-resistant strains highlights the need to identify novel drug targets (10).

Bacterial cell wall synthesis is a common target of many antibiotics. Considerable progress has been made to delineate the enzymatic pathways involved in peptidoglycan synthesis (11, 12), and our understanding of the mechanistic basis of cellular shape maintenance (13–15). In rod-shaped bacteria longitudinal and septal peptidoglycan synthesis are thought to be coordinated by two overlapping, cell wall-associated multi-protein enzymatic complexes. These complexes consist of various peptidoglycan-synthetic penicillin-binding proteins (PBPs) and associated proteins, along with cytoskeletal proteins, which together coordinate the synthesis of the sacculus and maintain cell shape. The bacterial actin homologue, MreB, is thought to direct longitudinal peptidoglycan synthesis, whilst FtsZ, a tubulin homologue, directs septal peptidoglycan synthesis and division (16–21). However, significant questions remain about the functions and interactions of many individual members of the peptidoglycan synthetic complexes.

Less-well studied factors in Gram-negative rod-shaped bacteria include the inner membrane proteins RodA and the penicillin binding protein 2 (PBP2). RodA is a member of the shape, elongation, division, sporulation (SEDS) family of proteins, and PBP2 is a class B penicillin binding protein (bPBP) and a major longitudinal cell wall transpeptidase. Its enzymatic activity is dependent upon RodA, whose function until recently remained unknown (22–25). Two important studies identified RodA as a peptidoglycan polymerase and this observation has led to a paradigm shift in our understanding of cell wall biogenesis (26, 27). In the new model proposed, peptidoglycan biogenesis involves two classes of peptidoglycan polymerases, class A penicillin binding proteins, and now also the SEDS-family protein RodA, partnering with PBP2 for transpeptidase cell wall cross-linking. These findings suggest that RodA may be a prime target for the next generation of antimicrobials to combat antibiotic-resistant bacterial pathogens, indeed a number of small molecules which target these enzymes have recently been identified (28). The inactivation of PBP2 or RodA in *E. coli* causes cells to form giant spheres, which eventually lyse. These phenotypes are almost identical to those seen with the inactivation of MreB, MreC, MreD or RodZ, other essential cell shape-determinant proteins which are also thought to form part of the longitudinal peptidoglycan synthase complex (25). The cytoskeletal MreB protein and associated MreC and MreD proteins appear to be linked to the enzymatic machinery (e.g. PBP2 and RodA) via RodZ. The functions of these proteins are closely linked and interdependent. For instance, isolated amino acid substitutions in the periplasmic domains of PBP2 and RodA restored rod-shape to *E. coli* Δ*rodZ* cells in a manner associated with MreB (29).

Whilst bacterial cell shape maintenance and cell wall synthesis is an active area of research, few studies have considered the potential relationship between cell wall synthesis and bacterial virulence(30–35). Recently Garcia-del Portillo and colleagues identified *Salmonella* PBP2 and PBP3 homologues and additional paralogues designated PBP2_SAL_ and PBP3_SAL_ sharing just over 60% amino acid sequence identity (36). Although loss of PBP2_SAL_ had no significant impact on virulence in the mouse model, additional data suggested PBP2_SAL_ could play a role in adaptation to the intracellular lifestyle of *Salmonella* by contributing to the formation of a second elongasome (37). Several major virulence determinants in pathogens such as *Salmonella* consist of cell wall-associated or wall-spanning protein complexes that interact directly with the peptidoglycan layer. Surprisingly, the roles of PBP2 and RodA in the assembly and function of these supramacromolecular virulence determinants have not been investigated. Changes to this peptidoglycan-binding, or the synthesis and structure of peptidoglycan itself, may significantly affect virulence determinant functionality (38, 39). For example, the specific interactions of the flagella P-ring with the peptidoglycan are thought to provide physical stability to the flagella rotor (40). Mutations in the peptidoglycan-binding regions of various bacterial secretion systems, including flagella and Type 3 secretion systems (T3SSs), are linked with a loss of function (41–43). Furthermore, several studies have suggested that the cytoskeletal members of the peptidoglycan-synthetic complexes directly control the localisation of proteins within the cell envelope (44, 45). There is a gap in our knowledge how the bacterial cytoskeleton proteins and the peptidoglycan sacculus itself, are involved in the assembly and function of important cell wall-spanning virulence organelles in *Salmonella*.

Two independent type-3 secretion systems (T3SSs) are important virulence factors in *Salmonella*. The SPI-1 and SPI-2 T3SS apparatuses and effector proteins are each encoded on separate genomic islands (SPI: *<u>S</u>almonella* <u>p</u>athogenicity island) (46). These secretion systems are cell wall-spanning multi-protein needle complexes which inject virulence effector proteins directly into the host cell cytoplasm (38, 46, 47). The SPI-1 T3SS initiates the initial invasion of gut epithelial cells; SPI-1 effector proteins induce uptake of the bacterium by reorganising host cell actin filaments and interfering with signal transduction. SPI-2 is expressed at a later stage of infection and is essential for intracellular survival within a modified phagosome, the “*Salmonella*-containing vacuole” (SCV) (38, 47, 48). SPI-2 T3SS effectors are thought to assist bacterial replication and induce maturation of the SCV (49). SPI-2 is also essential for the escape of *Salmonella* from infection foci and for its spread in the tissues, a key element of bacterial pathogenesis (50). Flagella, large complex cell wall-spanning organelles which are structurally related to T3SSs, have also been demonstrated to assist with virulence in *Salmonella* (51–54).

In this study we delineate the contribution of RodA and PBP2 to the biology and also virulence of *Salmonella.* We demonstrate their important roles in regulating the expression of the membrane spanning SPI-1 T3SS virulence factor. Notably, we highlight that loss of RodA or PBP2 leads to an attenuated phenotype *in vivo*, highlighting the importance of these proteins as a new target for antimicrobial drugs which not only impact cell wall biogenesis but also importantly attenuate virulence.

## Results

### The *pbpA* and *rodA* genes are members of the *mrd* operon in *Salmonella*

The overall aim of this study was to analyse the role and essentiality of the *pbpA* and *rodA* genes in the biology and virulence of *Salmonella*. The *mrd* operon is conserved across both pathogenic and non-pathogenic Gram-negative bacteria, consisting of five closely linked genes: *ybeB* (*rsfS*), *ybeA* (*rlmH*), *pbpA* (*mrdA*), *rodA* (*mrdB*) and *rlpA* (Fig 1A). *in silico* analyses of *Salmonella* genomic DNA using the online BDGP Fruitfly and Softberry BROM programmes, and the TranstermHP algorithm, provided evidence to support this predicted operon. The software identified putative promoter sequences a short distance upstream from *ybeB* in *Salmonella*, and highlight the lack of predicted transcriptional terminator sequences within the operon (55–57).The polycistronic nature of the mRNA was confirmed RT-PCR, and luciferase transcriptional reporter assays confirmed the presence of a true major *mrd* operon promoter upstream of *ybeB* in *Salmonella* (data not shown). The *mrd* operon is well conserved between *S.* Typhi and *S.* Typhimurium, with 99% sequence identity at the DNA level (BLASTn/megablast analyses; (58)).

**Figure 1.**
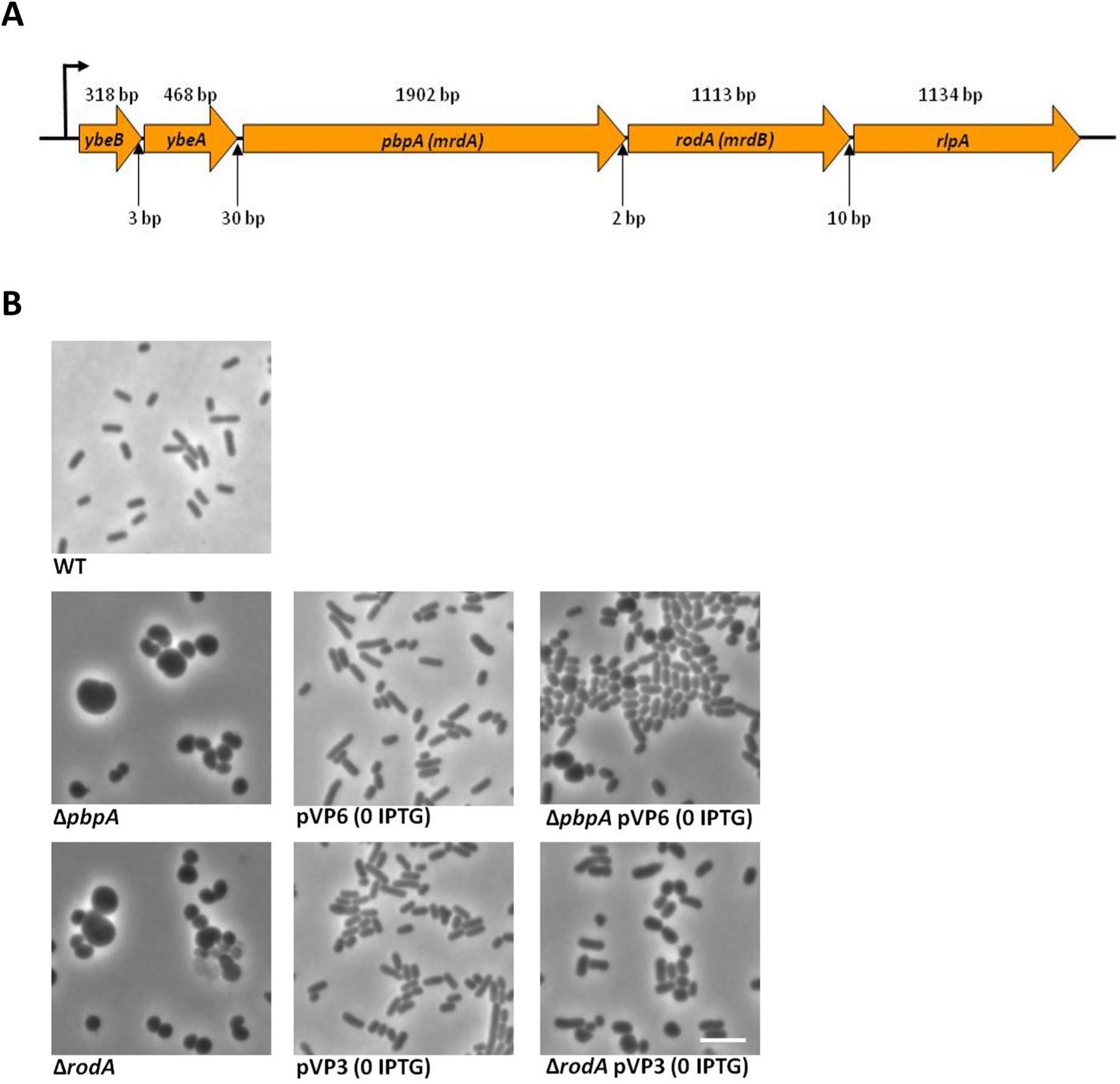
*pbpA* and *rodA* essential for rod-shape morphology. **(A):** Schematic Diagram of the putative *mrd* operon in *S*. Typhi. Position of putative operon promoter, gene sizes and inter-gene spacing as indicated. Operon drawn to scale. **(B):** Phase contrast images of *S.* Typhi BRD948 wild-type (WT), Δ*pbpA* and Δ*rodA* mutants, and complemented WT, Δ*pbpA* and Δ*rodA* cells. The latter express plasmid encoded YFP-fusion proteins of PBP2 and RodA respectively, as described in the text. Images taken of live cells at mid-log phase. All images to the same scale; scale bar indicates 5 µm.

### Construction of precise *pbpA* and *rodA* knockout *Salmonella* mutants and complementation of the mutant strains

Precise knockout mutants of both *pbpA* and *rodA* were generated in the *S.* Typhi BRD948 strain (59). Both Δ*pbpA* and Δ*rodA* mutants of *S.* Typhi BRD948 formed spherical cells which were prone to lysis (Fig 1B) (22, 25). The BRD948 Δ*pbpA* and Δ*rodA* mutants were subsequently complemented with plasmid-encoded N-terminal YFP-fusion proteins of either PBP2 (pVP6) or RodA (pVP3) from *E. coli* (60). The uninduced basal expression levels of PBP2-YFP and RodA-YFP in BRD948 Δ*pbpA* and BRD948 Δ*rodA* respectively were sufficient to restore rod-shape, demonstrating that the observed morphological defects resulted directly from the inactivation of *pbpA* or *rodA*. As a negative control, introduction of plasmid-encoded RodA-YFP into the Δ*pbpA* mutant did not complement the morphological defects (Fig 1B). Hence both PBP2 and RodA proteins are required to maintain rod-shape in *Salmonella* either directly, or through their relationship with other members of the peptidoglycan synthase machinery.

### Expression and secretion of SPI-1 and SPI-2 virulence effector proteins in *Salmonella* Δ*pbpA* and Δ*rodA* mutants

We systematically assessed the expression and functionality of the SPI-1 and SPI-2 T3SSs virulence determinants in *S.* Typhi Δ*pbpA* and Δ*rodA* mutants. In order to investigate SPI-1 and SPI-2 T3SS protein expression, assembly, and function, secreted protein fractions were isolated from the parent (WT), and isogenic Δ*pbpA* and Δ*rodA* BRD948 cultures, grown in either SPI-1-inducing or SPI-2-inducing conditions. These fractions were then analysed by SDS-PAGE and western blotting using antibodies against SPI-1specific effector proteins or against tagged SPI-2 effector proteins.

Whole cell lysates were analysed alongside secreted protein preparations, to assess both intracellular expression and T3SS-mediated secretion of the effector proteins (Fig 2 A-C). To study the ability of SPI-2 T3SS to secrete effector proteins, WT, Δ*pbpA* and Δ*rodA* BRD948 cells as well as *S.* Typhimurium Δ*ssaV* (SPI-2 non-secreting) strains were transformed with a plasmid-encoded haemagluttinin-tagged SseJ SPI-2 effector (pWSK-*sseJ*2HA) (38). SPI-2-mediated SseJ-2HA expression and secretion was detected on western blots using antibody specific to the haemagluttinin tag (Fig 2B).

**Figure 2:**
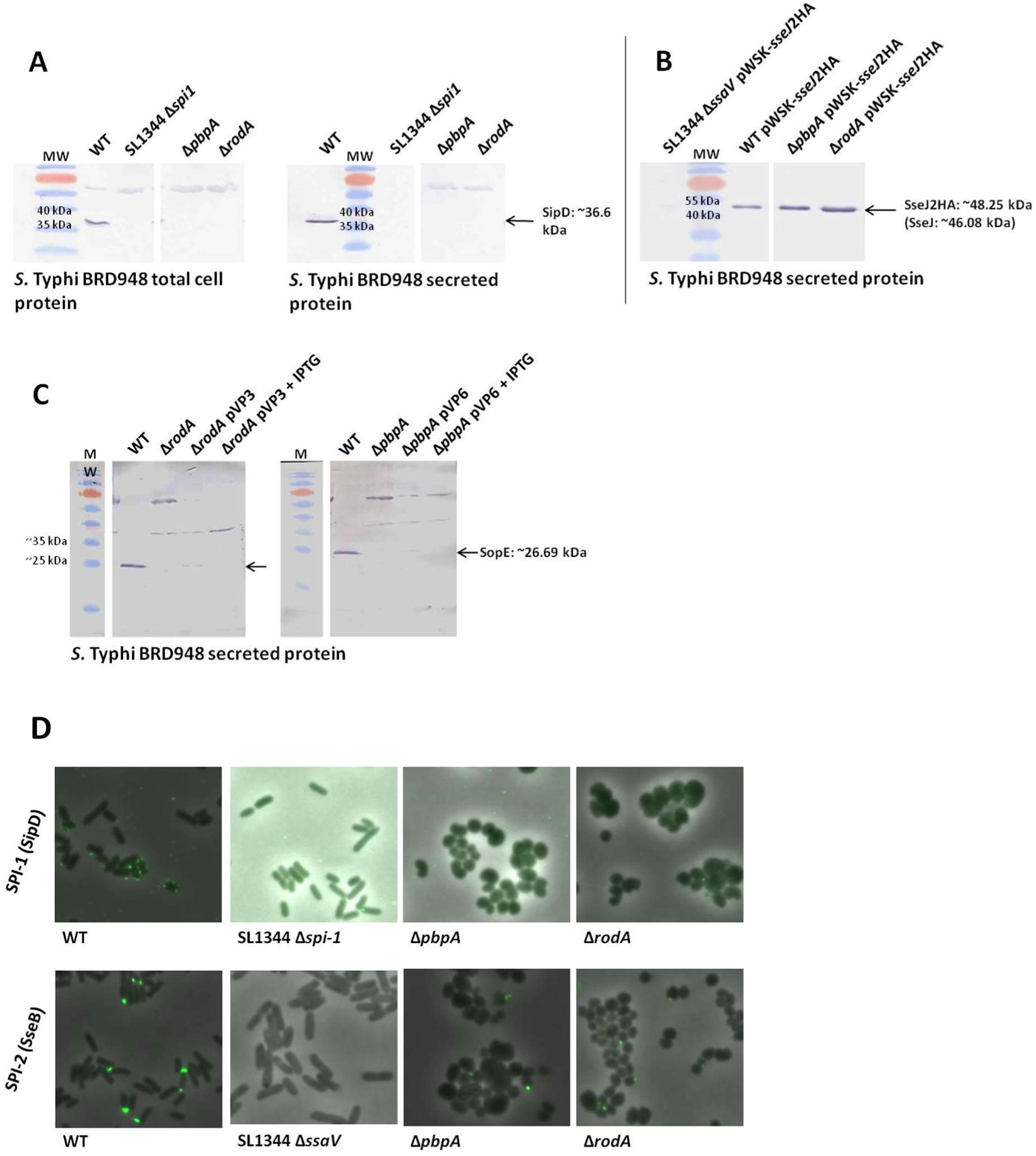
SPI-1 and SPI-2 T3SS effector protein expression in *Salmonella* (BRD948) isogenic parent, Δ*pbpA* and Δ*rodA* cells. **(A)** Western blots of total cell protein and secreted protein fractions of BRD948 parent and mutant cells, probed with antibodies against the SipD SPI-1 effector protein. *S*. Typhimurium SL1344 Δ*spi1* = negative control. **(B)** Western blots of secreted protein fractions of WT, Δ*pbpA* and Δ*rodA* cells expressing plasmid encoded HA-tagged SseJ (SPI-2 effector protein). Blot was probed with antibodies against the HA haemagluttinin tag. SL1344 Δ*ssaV* SPI-2 non-secretor = negative control. **(C)** Western blots of secreted protein fractions of WT/Δ*pbpA*/Δ*rodA* and complemented Δ*pbpA* and Δ*rodA* cells (+/− IPTG), probed with antibodies against the SopE SPI-1 effector protein. SPI-1 western blots were repeated on a number of occasions with a variety of SPI-1 effector protein-specific antibodies. **(D)** Immunofluorescence microscopy images overlaid with phase contrast images. Mid log-phase growth cells were fixed and probed with primary antibodies specific to SPI-1 or SPI-2 T3SS needle translocon proteins (SipD & SseB respectively), followed by fluorophore-conjugated secondary antibodies. The locations of individual T3SS needles are visualised as fluorescent green spots around the cell peripheries (SPI-1) or at the cell poles (SPI-2). SL1344 Δ*spi1* (SPI-1 non-secretor) and SL1344 Δ*ssaV* (SPI-2 non-secretor) strains were used as negative controls. All images to the same scale.

These assays demonstrated that SPI-1 effector proteins were neither secreted nor expressed intracellularly in either BRD948 Δ*pbpA* or Δ*rodA* cells, although effector protein expression and secretion was evident in the control WT strain and in complemented Δ*pbpA* and Δ*rodA* mutants expressing PBP2-YFP or RodA-YFP respectively (Fig 2A and 2C). In contrast, SseJ was visible in both the whole cell protein fraction and the secreted protein fractions of both BRD948 Δ*pbpA* and Δ*rodA*, demonstrating that SPI-2 effector proteins were expressed and secreted (Fig 2B).

To exclude the possibility that the presence of the SseJ protein in round-cell mutant secreted protein fractions may have been due to cell lysis, rather than active secretion through the SPI-2 needle, BRD948 WT, Δ*pbpA* or Δ*rodA* cells grown in SPI-2-inducing conditions were prepared for immunofluorescence microscopy. Samples were labelled with antibody specific to SseB, a SPI-2-secreted protein which forms part of the translocon at the outer tip of the T3SS needle (61). Immunofluorescence microscopy enabled the visualisation of cell surface SPI-2 needle complexes on both WT and round-cell mutant cells, but not at the surface of *S.* Typhimurium Δ*ssaV* cells (Fig 2D). This suggests that SPI-2 needles were correctly assembled and functional in BRD948 Δ*pbpA* and Δ*rodA* cells. By comparison, SPI-1 needles were not visible at the surface of BRD948 Δ*pbpA* and Δ*rodA* cells fixed and labelled with antibody specific to SipD, a secreted effector which forms part of the SPI-1 translocon (Fig 2D).

### Motility and Vi capsule expression in *S.* Typhi Δ*pbpA* and Δ*rodA* strains

The Vi (virulence) capsule is a key virulence factor, a polysaccharide which is produced by several bacteria in particular Typhi, Paratyphi C, *Citrobacter freundii* and *Salmonella* Dublin. Whilst the Vi capsule is immunogenic, infection can still occur without the polysaccharide (62). The expression of flagella and Vi capsule were determined phenotypically. The motility of both the *S*. Typhi Δ*pbpA* and Δ*rodA* mutant and complemented strains, was examined by measuring their migration through semi-solid agar. Δ*pbpA* and Δ*rodA S.* Typhi mutants were non-motile; ~60-80% motility was restored in complemented Δ*pbpA* pVP6 and Δ*rodA* pVP3 strains. (Fig 3A). Expression of the *S.* Typhi-specific Vi antigen capsule was examined in the Δ*pbpA* and Δ*rodA* mutants by means of slide agglutination assays using Vi-agglutinating serum. Clear agglutination in both WT and mutant BRD948 cells demonstrated that Vi capsule remained actively expressed (Fig 3D).

**Figure 3:**
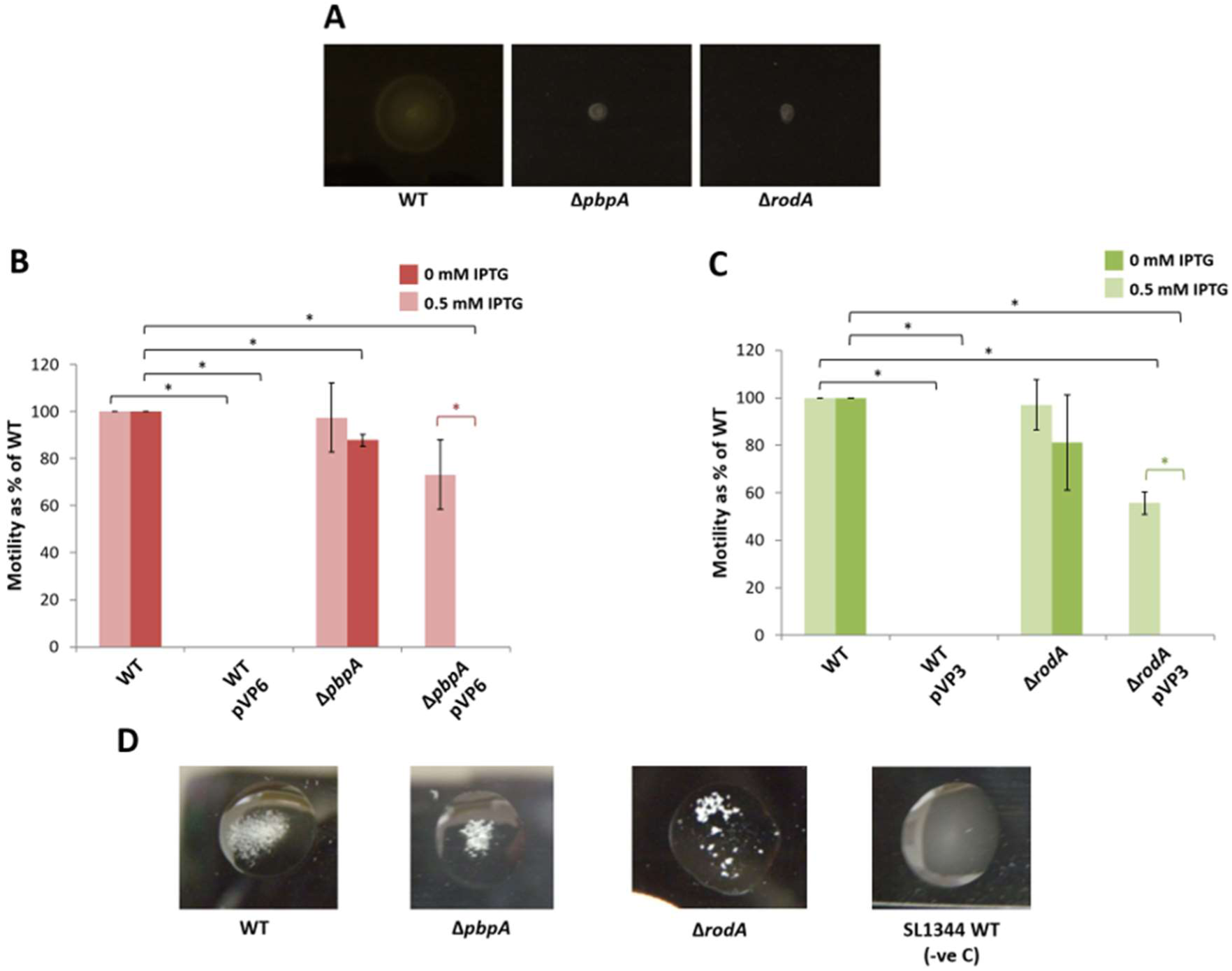
Motility assays of BRD948 WT, Δ*pbpA* and Δ*rodA* strains. **(A)** Representative image of motility assay plates showing growth and spread of WT and mutant cells through semi-solid agar. Motility was measured in terms of extent of spread (diameter) from site of inoculation. Motility assays were conducted for Δ*pbpA* and complemented Δ*pbpA* pVP6 strains +/− IPTG **(B)**, and for Δ*rodA* and complemented Δ*rodA* pVP3 strains +/− IPTG **(C).** Motility (i.e. diameter of growth) of the mutant strains was measured as a percentage of that of the WT. Assays were conducted a minimum of 3 times. Statistically significant differences are indicated by an asterisk. **(D)** Vi capsule expression in BRD 948 WT, Δ*pbpA* and Δ*rodA* cells. Images of slide agglutination assays, showing cell clumping upon addition of Vi-specific serum to liquid cultures. Clumping phenotype is indicative of Vi capsule expression. *S.* Typhimurium WT was used as negative control.

### SPI-1 and flagella virulence gene expression in *S.* Typhi Δ*pbpA* and Δ*rodA* mutants

SPI-1 and flagella gene expression were subsequently examined in the round-cell *S.* Typhi Δ*pbpA* and Δ*rodA* mutants with the aim of identifying the genetic basis of the observed SPI-1 and motility expression defects. Expression of the SPI-1 T3SS is regulated by and dependent upon three SPI-1-encoded transcriptional activators: HilA (the SPI-1 master regulator), HilC and HilD. HilA expression is controlled upstream by both HilC and HilD, although HilD appears to be more crucial since the majority of upstream environmental signals and global regulatory pathways seem to converge on HilD, post-transcriptionally regulating its expression (47, 54, 63–65).

In *Salmonella*, flagella gene expression is tightly regulated in a hierarchical manner, closely coordinated according to the successive stages of flagella assembly. At the peak of this hierarchy are FlhD and FlhC, which form a hetero-oligomer to transcriptionally regulate a global array of genes. *flhDC* expression is absolutely required for flagella protein expression and assembly, and hence motility (54, 66–68). Notably, the regulation of both SPI-1 and flagella gene expression is significantly inter-linked (54, 69, 70). Just downstream of FlhDC in this regulatory hierarchy is FliF, an integral flagellar basal body protein. *fliF* expression occurs very early on in during flagella assembly, FliF being the first flagella protein to be incorporated into the cell wall (66, 68, 71).

*hilD* and *fliF* were therefore ideal candidates through which to examine SPI-1 and flagella gene expression. The promoter regions of *hilD* and *fliF* were each individually cloned into the pMK1*lux* luciferase transcriptional reporter plasmid. pMK1*lux-hilD* and pMK1*lux-fliF* were subsequently introduced into WT, Δ*pbpA* and Δ*rodA* BRD948 strains. Transcriptional reporter assays were conducted, whereby the relative luminescence emitted from WT, Δ*pbpA* and Δ*rodA* cultures was measured as an indication of *hilD* or *fliF* promoter activity – and consequently of SPI-1 or flagella gene expression. Both *hilD* and *fliF* were strongly downregulated in *S.* Typhi Δ*pbpA* and Δ*rodA* mutants (Fig 4A and 4B). *flhDC* expression was also examined in these mutants using pRG38, an alternative luciferase transcriptional reporter plasmid harbouring the *flhDC* promoter region. Transcriptional reporter assays showed that *flhDC* was significantly downregulated in the Δ*rodA* strain, and almost completely downregulated in the Δ*pbpA* cells, compared to WT (Fig 4C).

**Figure 4:**
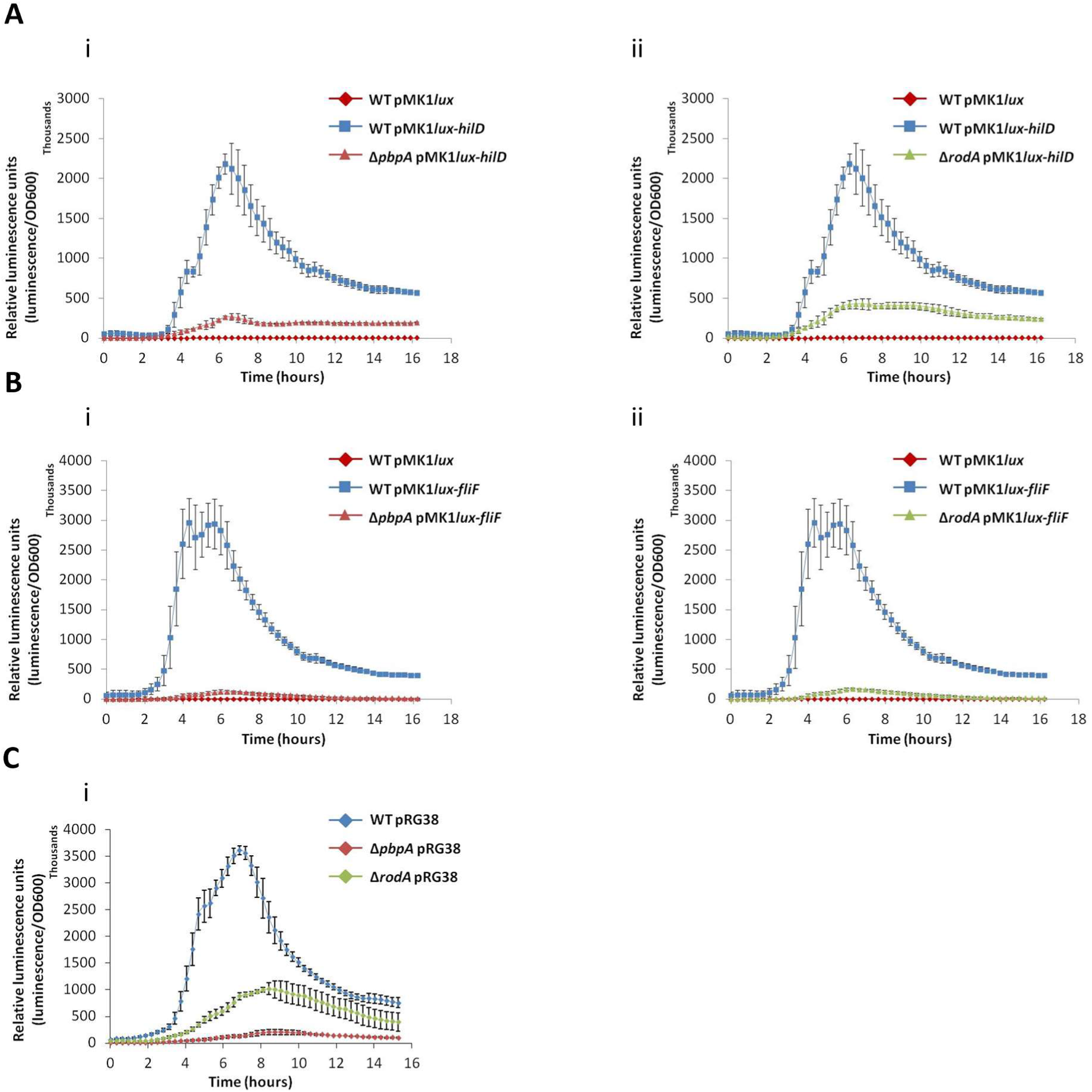
Flagella and SPI-1 T3SS gene expression in BRD948 isogenic parent (WT), Δ*pbpA* and Δ*rodA* strains. Transcriptional reporter assays were conducted to measure relative luminescence emitted from WT and mutant cells expressing the luminescence reporter plasmids pMK1*lux-hilD,* pMK1*lux-fliF* or pRG38, in which the *lux* operon is cloned just downstream of the *hilD*, *fliF* and *flhDC* promoter regions, respectively. Thus, light emission is representative of promoter gene activity and *lux* gene expression. Average measurements of three independent experiments are given. A) Strains harbouring pMK1*lux*-hilD i)ΔpbpA (red triangle) ii)ΔrodA (green triangle). B) Strains harbouring pMK1*lux*-fliF i)ΔpbpA (red triangle) ii)ΔrodA (green triangle). WT containing empty pMKlux indicated by a red diamond, WT containing pMK1*lux-gene* is indicated by a blue square. C) Strains harbouring the pRG38-flhDC plasmid: WT (blue), ΔpbpA (red) and ΔrodA (green).

### Complementation of the SPI-1 and motility defects in *S.* Typhimurium Δ*pbpA* and Δ*rodA* mutants

To investigate the roles of *rodA* and *pbp2* in virulence *in vivo*, we wished to utilise the well-established *S.* Typhimurium mouse model. This approach allowed us to conduct experiments in *S.* Typhimurium that mirrored those previously performed in *S.* Typhi, thereby facilitating a direct comparison of the impact of these genes on virulence *in vivo*. SPI-1 T3SS and flagella gene expression were also shown to be repressed in *S.* Typhimurium Δ*pbpA* and Δ*rodA* mutants. *hilD* expression was subsequently restored *in trans* in the *S.* Typhimurium SL1344 Δ*pbpA* and Δ*rodA* mutants by introducing plasmid-encoded *hilD*, expressed under the control of the arabinose-inducible P_BAD_ promoter (pBAD-*hilD*). The uninduced, basal level expression from pBAD-*hilD* was sufficient to restore effector protein expression in both SL1344 Δ*pbpA* and Δ*rodA* (Fig 5). Expression of *flhDC* was also restored *in trans* in SL1344 Δ*pbpA* and Δ*rodA* by transducing cells with a P22 phage lysate harbouring ‘TPA14’, a chromosomally-encoded vector in which *flhDC* expression is controlled by a tetracycline-inducible P*_tetA_* promoter (72). P22 phage is *S.* Typhimurium-specific, and so this vector could not be transduced into the *S.* Typhi strains (73). Motility assays demonstrated that SL1344 Δ*pbpA* and Δ*rodA* cells expressing TPA14 recovered significant motility in the absence of additional tetracycline (Fig 6). Thus expression of the HilD or FlhDC regulators was sufficient to restore the expression and assembly of functional SPI-1 needles or flagella, respectively, in Δ*pbpA* and Δ*rodA* mutants.

**Figure 5:**
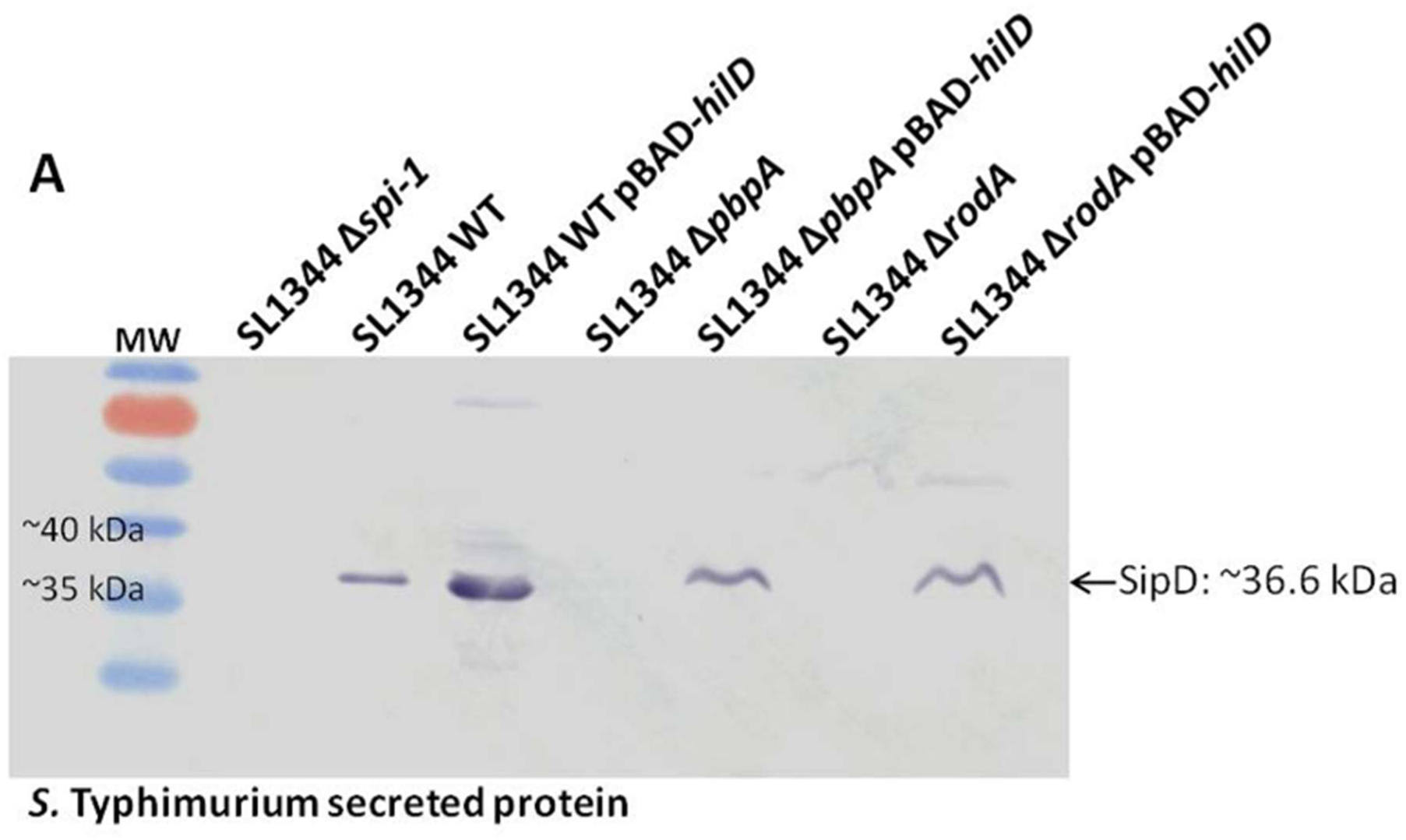
SPI-1 T3SS expression in *S.* Typhimurium (SL1344) isogenic parent, Δ*pbpA* and Δ*rodA*, and Δ*pbpA* and Δ*rodA* harbouring pBAD-*hilD*, in which the SPI-1 *hilD* regulator is expressed under the control of an arabinose-inducible promoter. Western blots of secreted protein fractions of WT and mutant cells, probed with antibodies against the SipD SPI-1 effector protein. *S*. Typhimurium SL1344 Δ*spi1* = negative control

**Figure 6:**
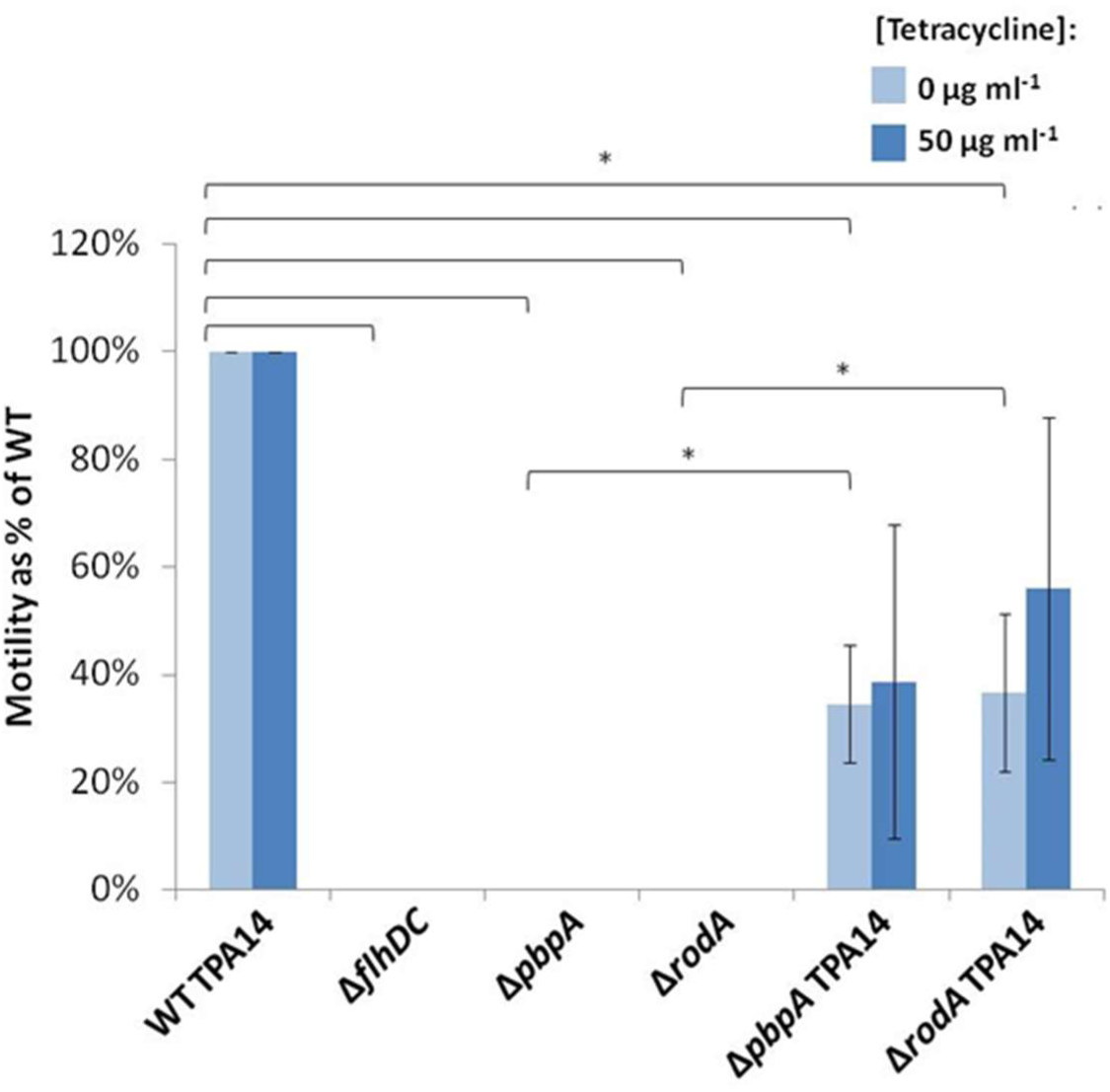
Motility assays of *S.* Typhimurium SL1344 WT, Δ*pbpA* and Δ*rodA*, and TPA14-complemented Δ*pbpA* and Δ*rodA* strains, grown in semi-solid agar +/− 50 µg/ml tetracycline. In strains harbouring TPA14, *flhDC* is expressed chromosomally under the control of the tetracycline-inducible P_tetA_ promoter. Motility (i.e. diameter of growth) of the mutant strains was measured as a percentage of that of the WT harbouring TPA14. Assays were conducted a minimum of 3 times. Statistically significant differences are indicated by an asterisk.

### Virulence of *S.* Typhimurium Δ*pbpA* and Δ*rodA* mutants *in vivo*

Given that SPI-2-T3SS expression and secretion function appeared to be largely unaffected in Δ*pbpA* and Δ*rodA* cells, we wished to investigate whether these strains remained virulent *in vivo*. In mice *S.* Typhimurium grows and spreads in the tissues, providing a model for systemic salmonellosis of humans and other animal species (74). WT, Δ*pbpA* and Δ*rodA* strains of SL1344 were inoculated into C57BL/6 mice intravenously rather than orally in order to bypass the SPI-1-dependent stages of infection as we have already eluded the SPI-1 T3SS is down-regulated in expression. Growth of the strains were determined by counting the numbers of viable bacteria in the spleens and livers, and onset of clinical symptoms were monitored over a 28 day period post-infection.

The wild-type strain grew at the expected rate of 10-fold per day and leading to clinical symptoms in under 5 days. However in contrast the Δ*pbpA* and Δ*rodA* strains were attenuated in their growth/survival *in viv*o, after an initial reduction in bacterial counts in the tissues, cell numbers increased slightly over a 6-day period, before steadily declining in both the liver and spleen as they were overcome by the host immunity (Fig 7).

**Figure 7:**
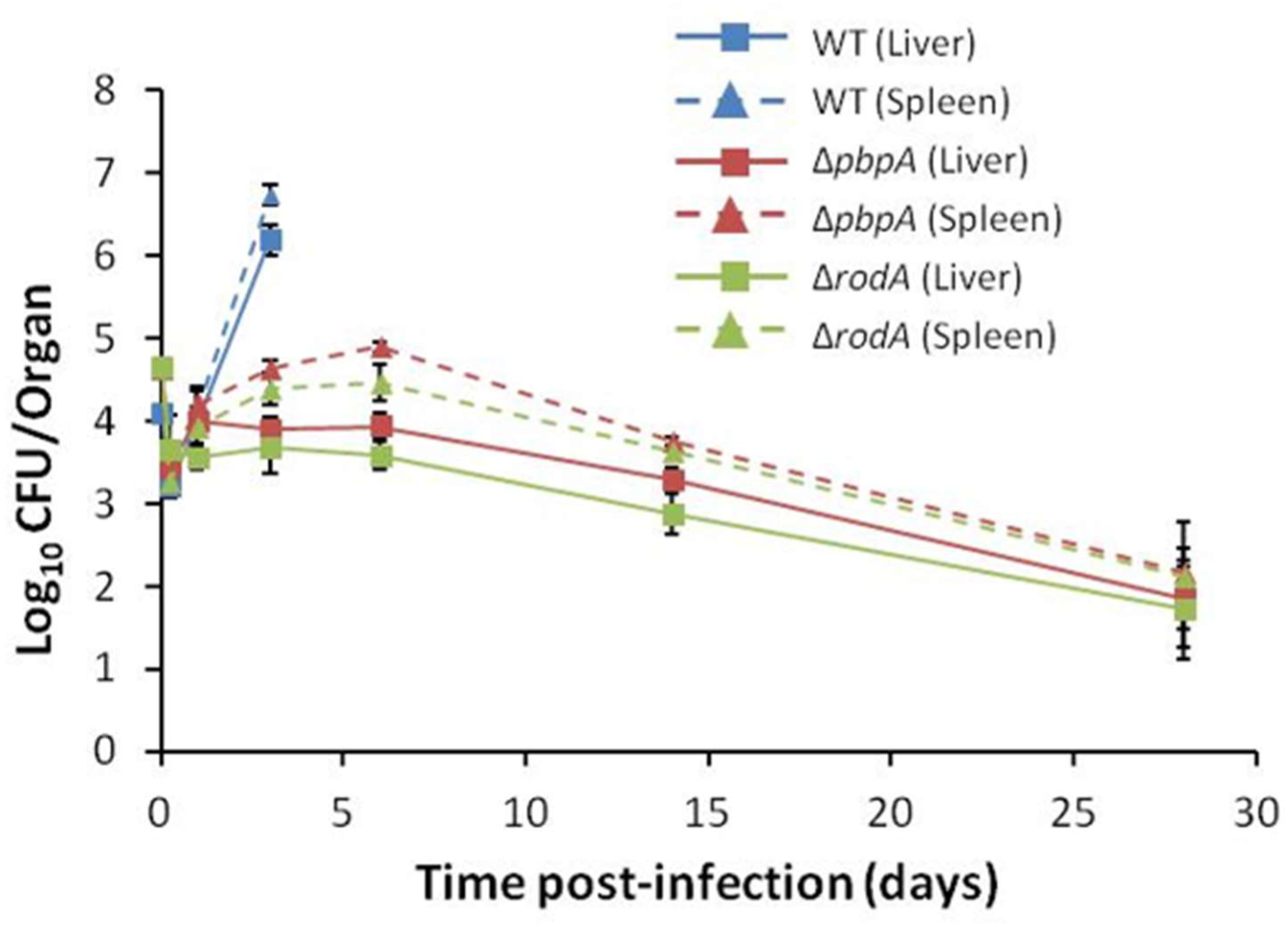
*in vivo* virulence assays of *S*. Typhimurium WT (SL1344), Δ*pbpA* and Δ*rodA* strains in mice, as a model for typhoid fever. For each strain, mice were inoculated intravenously with 10^4^ cells. Bacterial viable counts were subsequently performed at various time-points post-infection, to determine bacterial load in the liver and spleen. In the case of the WT strain, mice were euthanised at day 3. Average bacterial load values from 4 mice are shown (error bars = standard deviation).

### Global transposon mutagenesis screen to identify regulators of SPI-1 and motility in Δ*pbpA* and Δ*rodA* mutants of *S.* Typhimurium

We wished to uncover both the regulatory mechanisms and candidate transcriptional regulators involved in controlling SPI-1 T3SS and motility expression in response to the inactivation of *pbpA* or *rodA*. A transposon mutagenesis screen was therefore conducted to identify mutants in which flagella gene expression was restored due to the disruption of upstream regulators. This screen utilised a P22 phage lysate library, pooled from approximately 12000 Tn*10*d(Tc) individual insertion mutants.

In line with *S.* Typhi, pMK1*lux-fliF* reporter assays demonstrated that *fliF* was almost completely downregulated in *S.* Typhimurium SL1344 Δ*pbpA* and Δ*rodA* mutants (Fig 4B). SL1344 Δ*rodA* pMK1*lux-fliF* cultures were therefore transduced with the Tn*10*d(Tc) library. Approximately 7200 transductant colonies were then screened for recovery of luminescence, and hence restoration of flagella gene expression. The visible contrast in luminescence emission between the parent WT and Δ*rodA* strains greatly facilitated the rapid identification of Δ*rodA* pMK1*lux-fliF* transposon mutants with recovered luminescence, simply upon exposure to light-sensitive film (Figure 8A). The genomic position of each Tn*10*d(Tc) insertion was subsequently identified using arbitrary PCR and sequencing. In this way 14 positive luminescent revertants were identified, 13 of which harboured Tn*10*d(Tc) lesions at various points within the *rcsC* gene. In the remaining revertant Tn*10*d(Tc) was inserted within the 3’ end of the *pagN* gene (Figure 8B). In addition to recovered *fliF* expression, motility and SPI-1 effector protein expression were also restored to differing extents in several of these mutants (Figure 9A, B). However, SPI-1 needles were not visible at the cell surface, suggesting that SPI-1 needle assembly and secretion was still disrupted (Figure 9C).

**Figure 8:**
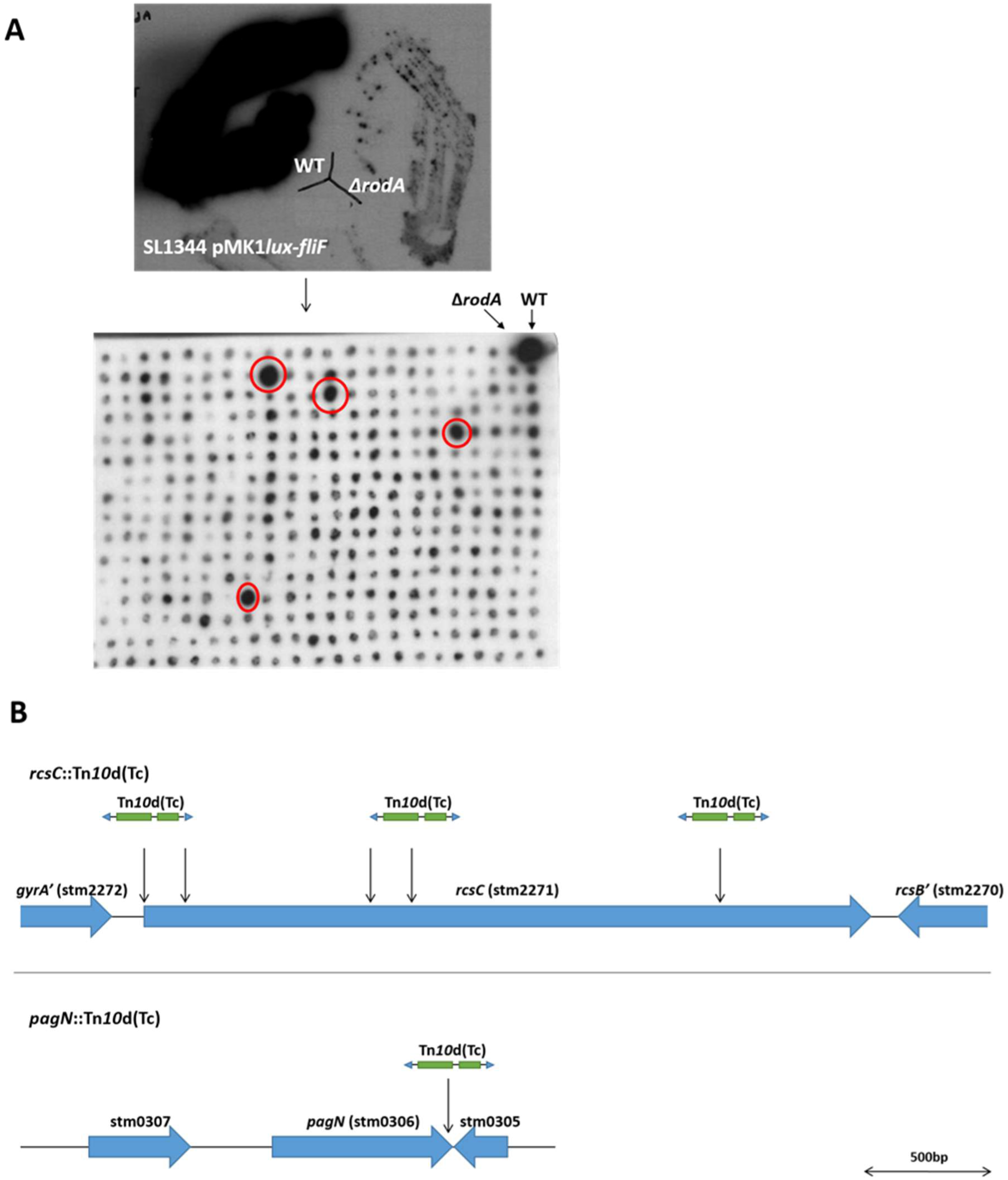
Development of a genome-wide Tn*10*d(Tc) transposon mutagenesis screen to identify mutants with recovered flagella-gene expression. (A) The contrast in luminescence emitted from WT and Δ*rodA* cells expressing the pMK1*lux-fliF* reporter plasmid was both stark and readily distinguishable by blotting plates of fresh cultures onto filter paper and exposing them to light-sensitive film. This phenotype was utilised in the development of a mutagenesis screen, whereby individual SL1344 Δ*rodA* pMK1*lux-fliF* Tn*10*d(Tc) mutant colonies were systematically screened for recovered luminescence, indicative of restored *fliF* expression. **(B)** Schematic diagrams demonstrating the genomic positions of Tn*10*d(Tc) mini-transposon insertions in relevant luminescent revertant strains of SL1344 Δ*rodA* pMK1*lux-fliF* (indicated by arrows). Tn*10*d(Tc) mini-transposons not drawn to scale.

**Figure 9:**
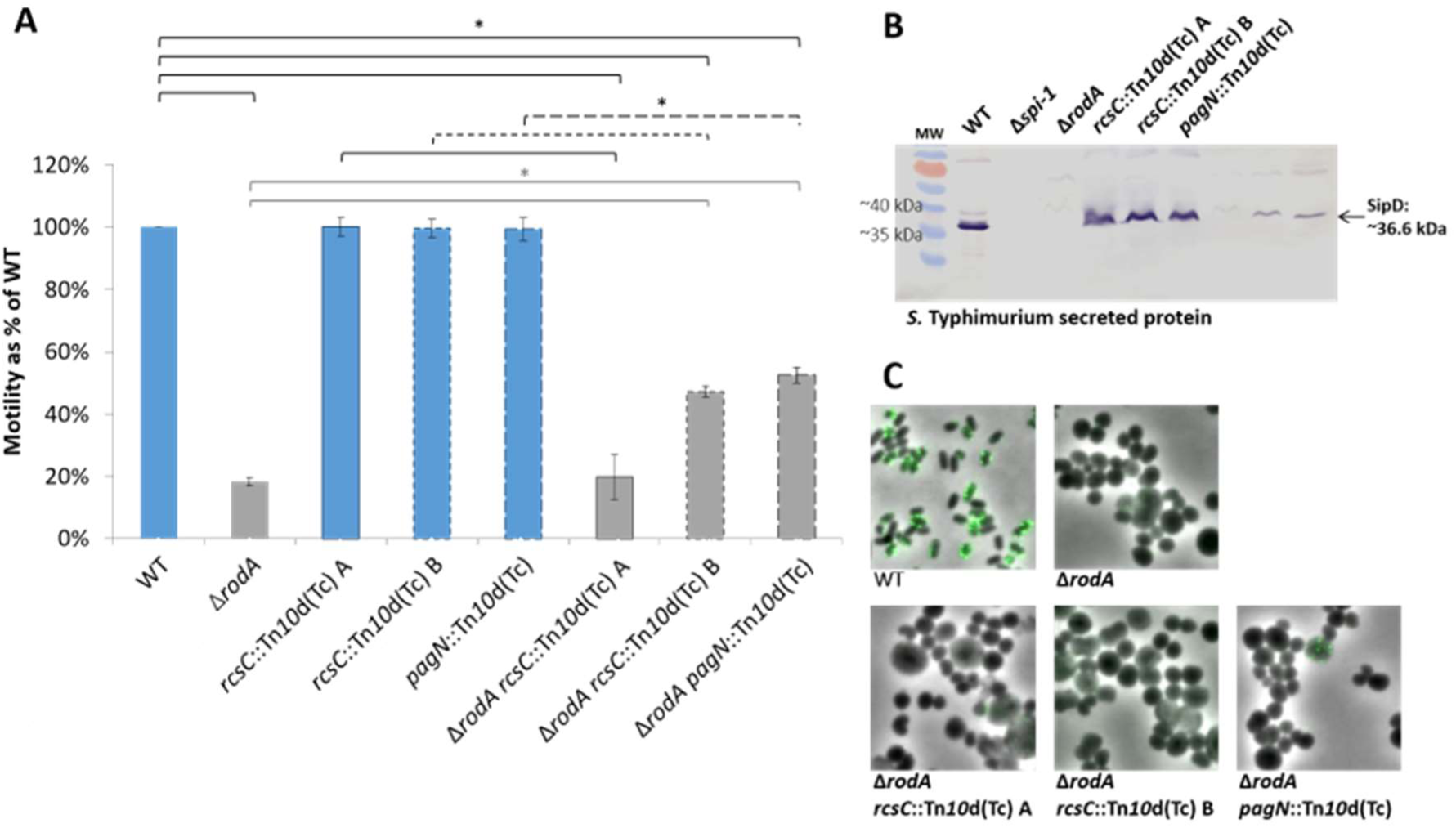
Phenotypic analysis of *S.* Typhimurium *rcsC*::Tn*10*d(Tc) and *pagN*::Tn*10*d(Tc) insertional mutants. Both *rcsC*::Tn*10*d(Tc) and *pagN*::Tn*10*d(Tc) mutants were transduced into the parent WT and Δ*rodA* strains, prior to further phenotypic analysis. (A) Motility assays of *S.* Typhimurium WT (blue bars), Δ*rodA* (green bars), +/− *rcsC*::Tn*10*d(Tc) or *pagN*::Tn*10*d(Tc) insertional mutations. Motility (i.e. diameter of growth) of both the Δ*rodA* and Tn*10*d(Tc) mutant strains was measured as a percentage of that of the WT parent. Assays were conducted a minimum of 3 times. Statistically significant differences are indicated by an asterisk. **(B)** Western blot of secreted protein fraction and **(C)** immunofluorescence microscopy images of WT, Δ*rodA* and Tn*10*d(Tc) mutant cells probed with antibodies against the SipD SPI-1 effector protein, showing effector protein secretion and cell surface T3SS needle expression respectively. *S*. Typhimurium SL1344 Δ*spi1* = negative control (not shown in C).

### Further analysis of RcsC and PagN as candidate regulators of virulence in response to *pbpA* or *rodA* inactivation

The role of the two candidate regulators was investigated further through the generation of specific non-polar knockout mutants of Δ*rcsC* or Δ*pagN* in both SL1344 and BRD948 Δ*pbpA* and Δ*rodA* strains, although attempts to generate a BRD948 Δ*pbpA* Δ*pagN* mutant were unsuccessful. Results from subsequent motility assays and western blot analyses to investigate SPI-1 effector protein expression, however, were inconclusive. Low but significant levels of motility were restored only in SL1344 Δ*pbpA* Δ*rcsC* double mutants (compared to the Δ*pbpA* parent strain), and not in any of the BRD948 double mutants (Figure 10). Low levels of SPI-1 effector protein expression were also seen only in the BRD948 Δ*rodA* Δ*pagN* mutant, although no SPI-1 needles were observed at the cell surface (Figure 11). In Δ*pagN* and Δ*rcsC* single mutants of *S.* Typhi and Typhimurium, SPI-1 secretion was not significantly affected; motility was very slightly reduced in *S.* Typhimurium Δ*pagN* (Figure 10).

**Figure 10:**
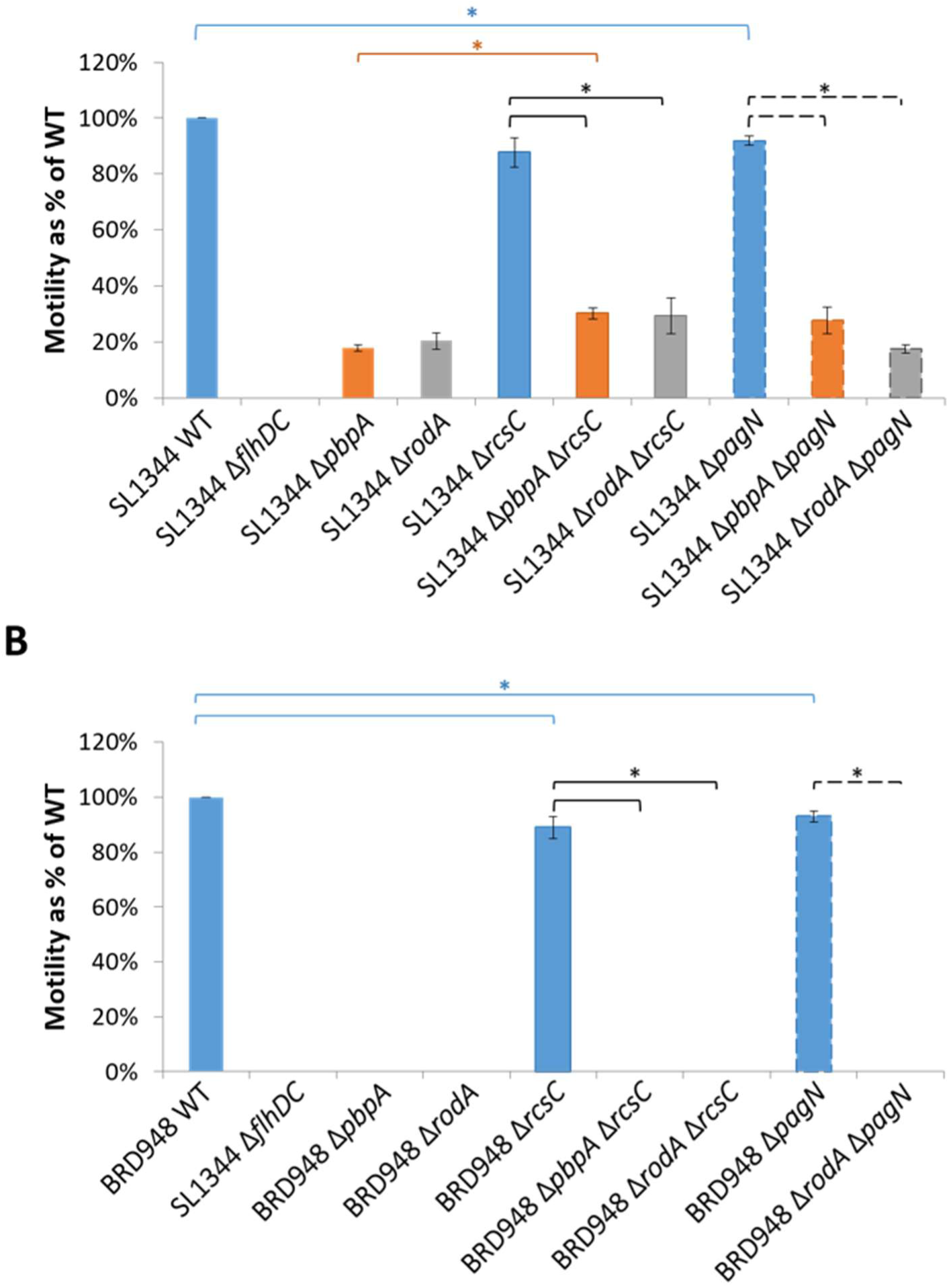
Motility assays of BRD 948/SL1344 WT, ΔpbpA, ΔrodA, ΔrcsC, ΔpagN single mutants, and ΔpbpA ΔrcsC/ΔpagN and ΔrodA ΔrcsC/ΔpagN double mutants, grown in semi-solid agar. Complete knockouts of ΔrcsC and ΔpagN were generated in both S. Typhimurium (A) and S. Typhi (B) WT, ΔpbpA and ΔrodA parent strains. Motility (i.e. diameter of growth) of each mutant was subsequently measured as a percentage of that of the respective WT parent. Assays were conducted a minimum of 3 times. Statistically significant differences are indicated by an asterisk.

**Figure 11:**
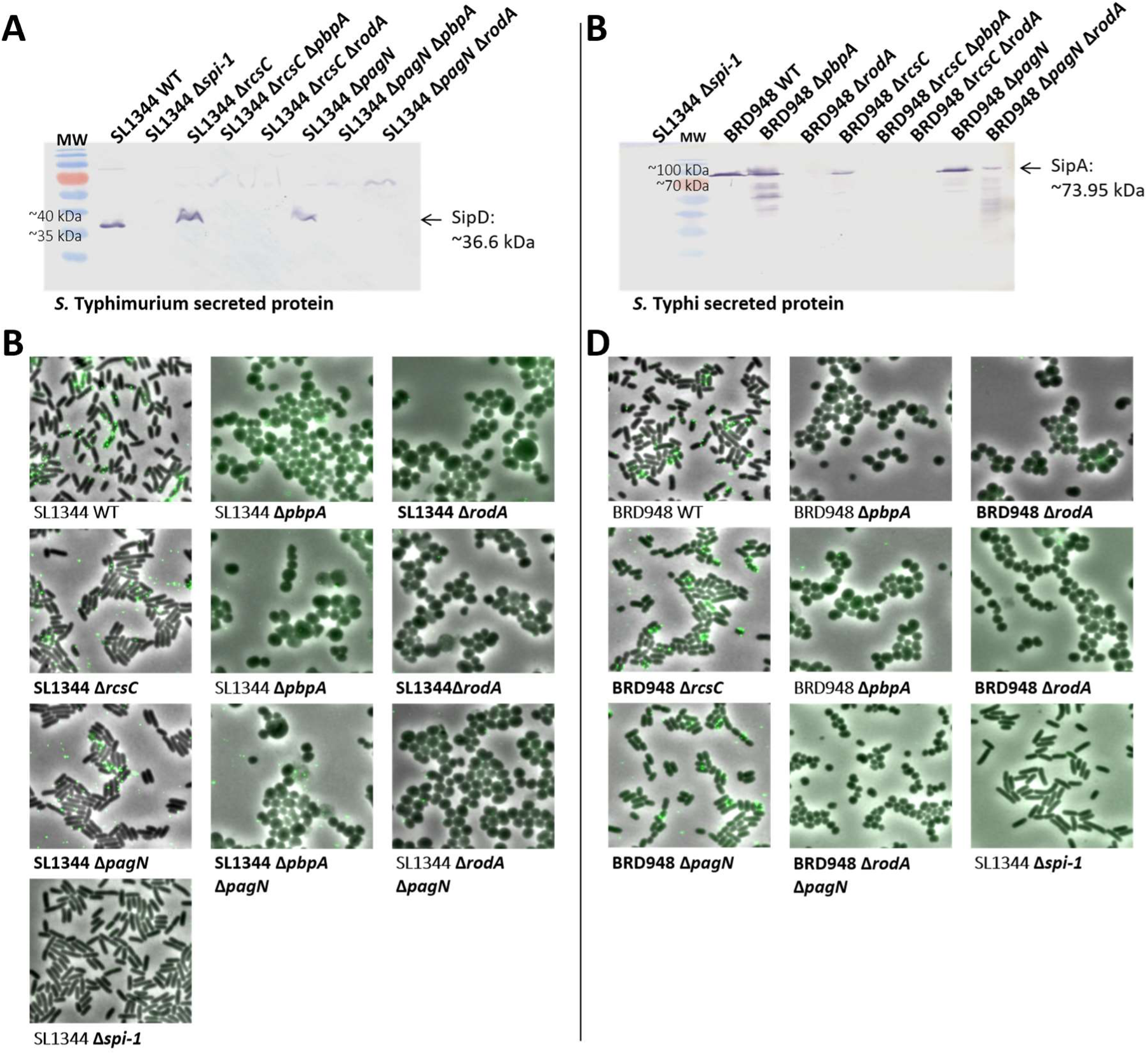
SPI-1 effector protein secretion/cell surface expression in Δ*rcsC* and Δ*pagN* mutants of WT, Δ*pbpA* and Δ*rodA*, in *S.* Typhimurium (A & C) and *S.* Typhi (B & D). A & C – Western blots of *S.* Typhimurium/*S.* Typhi secreted protein fractions of WT and mutant cells, probed with antibodies against the SipD or SipA SPI-1 effector proteins. *S*. Typhimurium SL1344 Δ*spi1* = negative control. B – Immunofluorescence/phase contrast microscopy images of mid log-phase growth *S*. Typhimurium/Typhi cells fixed and probed with primary antibodies specific to SipD, followed by fluorophore-conjugated secondary antibodies. The locations of individual T3SS needles are visualised as fluorescent green spots around the cell peripheries. SL1344 Δ*spi1* (SPI-1 non-secretor) was used as a negative control. All images are the same scale.

These conflicting results suggest that the regulation of virulence in Δ*pbpA* and Δ*rodA* mutants of *S.* Typhimurium, and perhaps particularly *S.* Typhi, is more complex and possibly multifactorial, involving input from a number of regulators. Further research is needed to clarify the role(s) of both transcriptional and post-transcriptional regulators in *Salmonella* round-cell mutants.

## Discussion

In rod-shaped bacteria such as *Salmonella*, longitudinal peptidoglycan synthesis is coordinated by multiprotein complexes comprising the peptidoglycan synthetic and transpeptidation enzymes, and associated cell shape-determinant proteins (16, 17, 20, 25, 31). Whilst PBPs and RodA are both required to catalyse peptidoglycan polymerization, the cytoskeletal proteins are thought to physically support and position the complexes within the bacterial envelope (22, 25, 29, 75, 76).

Only a small number of studies have investigated the relationship(s) between the peptidoglycan cell wall and bacterial pathogenesis (30–35). Specifically our understanding of how the functions of major cell wall-associated virulence determinants are affected as a result of perturbed cell wall synthesis have not been investigated in detail, though previous work from our lab demonstrated such relationships in *mre* mutants of *S.* Typhimurium (77).

RodA and PBP2 are both required to catalyse peptidoglycan polymerization and transpeptidation, The role of RodA makes it an attractive new target for the next generation of antibiotics. This study focused on PBP2 and RodA, proteins which are encoded contiguously within the *mrd* operon in Gram-negative bacteria such as *Salmonella* and *E. coli* (23). Complete knockout mutants of the *pbpA* and *rodA* genes were generated in *Salmonella enterica* serovar Typhi (*S.* Typhi), and shown to be non-polar on the expression of downstream genes. As was previously demonstrated in *E. coli* (25) and *Bacillus subtilis* (78, 79), *S.* Typhi and *S.* Typhimurium Δ*pbpA* and Δ*rodA* mutants lost their rod-shaped morphology. This phenotype was complementable upon expression of *E. coli* PBP2-YFP or RodA-YFP protein fusions *in vitro* (60).

In *S.* Typhi Δ*pbpA* and Δ*rodA* cultures grown in LB medium, significant heterogeneity in cell size and extensive cell lysis were both evident. By supplementing the culture medium with 20mM MgCl_2_ or CaCl_2_, Δ*pbpA* and Δ*rodA* cells stably propagated as small spheres with significantly reduced lysis. Similar phenotypes have also previously been observed in round-cell mutants of *Bacillus subtilis*, whereby cells exhibit some dependence on divalent cations for normal growth and morphology (80–82), possibly due to some effect on ‘stiffening’ of the cell envelope (80). However, the effects of magnesium or calcium in *B. subtilis*, a Gram-positive organism, may be somewhat dissimilar to *Salmonella*. Furthermore, the addition of MgCl_2_ or CaCl_2_ was sufficient to restore rod-shape in *B. subtilis*, but not in *Salmonella* round-cell mutants (Supplementary ?0 (80).

Increased expression of the essential cell division protein FtsZ is known to aid survival in round-cell mutants, potentially through facilitating the formation of complete Z-rings for cell division (25, 83). The addition of MgCl_2_/CaCl_2_ may prevent cell lysis in Δ*pbpA* and Δ*rodA* mutants either by indirectly activating *ftsZ* expression, through modulating production or activity of the alarmone ppGpp (84–87), which is known to regulate *ftsQAZ* transcription (88), or by directly promoting FtsZ oligomerisation and Z-ring stability (89). Increased ppGpp levels afford improved cell division and survival in *mrd* and *mre* mutants of *E. coli*, something attributed to increased FtsZ production (25, 90). Divalent cations are thought to physically interact with the lipopolysaccharide (LPS) layer in Gram-negative bacteria, bridging LPS molecules and strengthening both the cell envelope structure and barrier function (91, 92). However, high magnesium or calcium levels can also disrupt the Gram-negative outer membrane (93). Thus increased MgCl_2_/CaCl_2_ may help prevent cell lysis in round-cell mutants such as Δ*pbpA* and Δ*rodA* through a strengthening of the outer membrane.

Both *S.* Typhi Δ*pbpA* and Δ*rodA* mutants exhibited sensitivity to low temperatures, with reduced growth rates and significant cell lysis at 30°C. This observation was contrary to previous studies in *E. coli*, which showed that growth of *mre* and *mrd* operon mutants was improved in conditions permitting slow growth; i.e. at lower temperatures and in minimal media (25). However, a more recent study found that *E. coli* Δ*rodZ* mutants were unable to form colonies on LB agar at ≤30°C (75). It was proposed that the temperature sensitivity of the Δ*rodZ* mutants was due to a reduction in the quantity of peptidoglycan and disruptions to the overall cell wall integrity (29). This may also be the case for Δ*pbpA* and Δ*rodA* mutants of *Salmonella*.

By means of a range of phenotypic assays, it was demonstrated that virulence factor expression and functionality were significantly affected in Δ*pbpA* and Δ*rodA* mutants of *Salmonella*. As was previously observed in *S.* Typhimurium Δ*mreC* (77), both motility and the SPI-1 T3SS, important virulence factors in the initial stages of *Salmonella* infection, were almost completely downregulated in *S.* Typhi and Typhimurium Δ*pbpA* and Δ*rodA* cells from the level of the master regulators, *flhDC* and *hilD* respectively. To our surprise, under SPI-2-inducing conditions - which mimic the intracellular macrophage environment - the SPI-2 T3SS was both actively expressed in the Δ*pbpA* and Δ*rodA* mutants, and apparently functional; SPI-2 T3SS needles were observed at the cell surface of these mutants. Expression of the Vi capsule also remained fully active in *S.* Typhi Δ*pbpA* and Δ*rodA* cells.

Studies have revealed increased sensitivity to various external stresses in round-cell mutants (94, 95). Furthermore, cells treated with mecillinam, an antibiotic which specifically inactivates PBP2, showed a 50% reduction in the rate of peptidoglycan synthesis (22), and Δ*rodZ* mutants were shown to have reduced cell wall peptidoglycan content (31). These suggest that the cell envelope integrity and barrier function are significantly compromised in such cells. The peptidoglycan layer itself may function as a scaffold to support cell wall-spanning organelles; physical associations with peptidoglycan are known to be important for both flagella and T3SS function (39, 41–43). It was therefore expected that any perturbations to the peptidoglycan structure resulting from the inactivation of *pbpA* and *rodA*, could render the cells unable to assemble functional T3SSs or flagella.

In round-cell mutants such as Δ*pbpA* or Δ*rodA* mutants longitudinal cell wall synthesis is disabled; cell wall synthesis and cell growth seem to become dependent upon septal peptidoglycan synthesis, coordinated by FtsZ and associated proteins (22). SPI-2 needles are known to localise solely at the cell poles, whilst SPI-1 needles and flagella are expressed in multiple copies across the lateral surface of the cell (Figure 2) (47). It was therefore possible that SPI-2 T3SSs remained functional in Δ*pbpA* and Δ*rodA* mutants due to their polar localisation, within the ‘unaffected’ septal peptidoglycan regions.

However, upon restoring expression of the *hilD* and *flhDC* regulators *in trans* in *S.* Typhimurium Δ*pbpA* and Δ*rodA* cells, significant levels of SPI-1 effector protein expression and motility were restored, respectively. Furthermore, SPI-1 needles were observed at the cell surface in mutants expressing pBAD-*hilD*. It is possible that the reduced expression of these systems in Δ*pbpA* and Δ*rodA* cells with recovered *hilD*/*flhD* expression was due to structural defects within the cell wall. However, that the cell envelope in Δ*pbpA* and Δ*rodA* mutants remains sufficiently robust to support functional SPI-1 and flagella organelles is remarkable – particularly given the inherent structural defects. Overall, these results suggest that the observed SPI-1 and motility defects were due to specific downregulation of the two systems rather than an innate inability to sustain the assembly and functionality of the SPI- and flagella organelles.

In 2010, Nibe et al. described *E. coli* Δ*rodZ* in which Class 2 and 3 flagella genes, though not *flhDC*, were downregulated. Cells were non-motile but expressed flagella at the cell surface. They also isolated revertant cells in which flagella gene expression and motility were recovered. Interestingly, some wild-type morphology was also restored in these cells (31).

Virulence gene regulation and the stress responses are intrinsically linked in *Salmonella*. A number of global stress regulators, including two-component systems such as PhoP/PhoQ and OmpR/EnvZ, and particularly those involved in responses to envelope stress, also constitute major virulence gene regulators. These respond to environmental signals including osmolarity, nutrient availability and pH, controlling expression of a wide array of genes to promote cell survival and virulence (96–100). During an infection *Salmonella* are therefore able to sense their locality and express the appropriate virulence genes. Broadly-speaking, the invasion-associated SPI-1 and flagella genes are expressed early on *in vivo*, in response to the nutrient-rich gut lumen environment, whilst the SPI-2 system is expressed post-Invasion, in response to signals indicative of an intracellular environment. Expression of SPI-1 T3S and motility are therefore predominantly regulated differentially to SPI-2 genes, at least *in vitro* (97, 101–104).

We hypothesised that the observed downregulation of the invasion-associated SPI-1 and motility genes in the Δ*pbpA* and Δ*rodA* mutants, along with the maintenance of SPI-2 expression, resulted from the direct action of specific global stress/virulence regulators activated in response to perturbations to the cell wall’s integrity or barrier function. In support of this hypothesis, it was previously noted that reduced flagella expression in Δ*rodZ* mutants could be caused by stress signals activated through cell wall perturbations (31).

Our ongoing work sought to identify the regulator(s) involved, in order to better understand the more global impact of the Δ*pbpA* and Δ*rodA* mutations on *Salmonella* pathogenesis. No assumptions were made concerning the identity of these regulators. The Tn*10*d(Tc) mutagenesis screen approach therefore hoped to isolate both the most important and potentially previously unrecognised regulator(s). Two interesting candidates were identified – RcsC, and PagN. In both *S.* Typhimurium Δ*rodA rcsC*::Tn*10*d(Tc) and *pagN*::Tn*10*d(Tc) mutants motility and SPI-1 protein expression were restored. All mutants retained a spherical morphology, demonstrating that the motility and SPI-1 phenotypes were not due to recovery of RodA and/or PBP2 expression.

Gene function was likely disrupted in all *rcsC*::Tn*10*d(Tc)/*pagN*::Tn*10*d(Tc) revertants, since the Tn*10*d(Tc) insertions were all within the *rcsC* or *pagN* coding sequences. Furthermore, the Tn*10*d(Tc) positions varied between different *rcsC*::Tn*10*d(Tc) revertants, demonstrating that the revertant phenotype was not specific to the inactivation of a particular portion of *rcsC*. However, the recovery of significant motility and SPI-1 expression was not as clearly evident in Δ*pbpA* and Δ*rodA* Δ*rcsC*/Δ*pagN* precise double knockouts, of either *S.* Typhi or *S.* Typhimurium. The regulatory pathways involved are likely complex and multi-factorial, such that the inactivation of *rcsC* or *pagN* alone may be insufficient to fully rescue motility/SPI-1 expression or functionality. Differences in the nature of individual mutations may also affect the observed phenotypes.

The Rcs Phosphorelay is a complex and well-studied global envelope stress response pathway which regulates a large cohort of genes associated with survival in unfavourable conditions and maintenance of cell wall integrity (105–107). In addition to upregulating the colanic acid biosynthesis, SPI-2 and Vi biosynthesis genes, this system downregulates motility and SPI-1 in *Salmonella* (105, 108–111). The Rcs phosphorelay is the only pathway known to respond specifically to peptidoglycan-associated stress (30, 34). Thus RcsC represents a very likely candidate for playing a major role in virulence gene regulation in response to the inactivation of *pbpA* or *rodA*.

Several previous studies have pointed to a particular link between perturbations to the cell wall structure and associated PBP proteins, and activation of the Rcs phosphorelay. For instance, a study by Laubacher & Ades (30) demonstrated that the mecillinam-mediated inactivation of PBP2 in *E. coli* induced *rcs* gene expression.

Furthermore, a more recent study in *E. coli* isolated a mutant in a specific set of non-essential PBPs (PBP4, PBP5, PBP7 and AmpH), in which motility was completely downregulated due to the activation of the Rcs phosphorelay and also the CpxAR system, another regulator of envelope stress (33). Wild-type motility was restored with the overexpression of *flhDC* or the inactivation of RcsB or CpxAR, whilst some motility was recovered with the loss of RcsC. However, the Rcs-mediated response in this mutant was dependent on CpxAR.

Interestingly neither RcsB nor CpxAR were isolated as being important for the recovery of motility in *Salmonella* Δ*pbpA* or Δ*rodA*. In fact, although RcsC, the Rcs phosphorelay sensor-kinase, was isolated a number of times in the Tn*10*d(Tc) screen, no other major members of this two-component system were isolated – including the response-regulator RcsB, the intermediate RcsD protein, and the outer membrane activator RcsF, which is thought to detect peptidoglycan- or outer membrane-associated stress (112, 113). Likewise, *S.* Typhimurium Δ*mreC* Δ*rcsC* double mutants had significant recovery of both motility and SPI-1 protein expression. However, *rcsB, rcsD, rcsF* and *cpxAR* inactivation did not rescue SPI-1/motility expression in Δ*mreC* cells (77).

The RcsC protein possesses both kinase and phosphatase activity (114, 115). Unlike the other Rcs proteins, the inactivation of RcsC does not completely disrupt Rcs phosphorelay signalling; the deletion of *rcsB, rcsD* or *rcsF*, but not *rcsC* restored motility in motility-impaired *E. coli* mutants and RcsB could not be de-phosphorylated in Δ*rcsC* mutants (116). In a separate study, point mutations within *rcsC* caused increased RcsC expression (117). Akin to these findings, it may be that some residual activity of RcsC is required in round-cell mutants for the recovery of invasion-associated virulence gene expression – something that could be true of *rcsC*::Tn*10*d(Tc) but not Δ*rcsC* knockouts of *Salmonella* Δ*rodA* mutants. It is also possible that the individual phenotypic effects of specific PBP mutations affect the nature of the envelope stress response; thus other Rcs members, or the CpxAR pathway appear to be important for virulence gene regulation in some PBP mutants but not in Δ*pbpA*/Δ*rodA*/Δ*mreC* strains.

Several studies have independently identified an important role for the Rcs phosphorelay in the response to perturbed cell wall synthesis (30, 33). The function and impact of the Rcs phosphorelay signalling on both the general cell physiology and virulence gene regulation, is of considerable interest. Although the pathway can be activated by peptidoglycan stress via RcsF, currently no members of the Rcs regulon are known to be involved in peptidoglycan synthesis (30, 33, 34). It may be that the activation of major known members of the regulon compensates for the peptidoglycan synthesis defects. For instance, overexpression of the *ftsQAZ* genes is required for survival in round-cell mutants, something that was considered to result from the selection for secondary mutations. However, the Rcs phosphorelay is known to activate *ftsQAZ* expression (107); consequently *ftsQAZ*– overexpression may be a specific Rcs phosphorelay-regulated response in round-cell mutants. The Rcs phosphorelay may also activate an as yet unrecognised cohort of genes involved in peptidoglycan synthesis in round-cell mutants. In *E. coli* cells depleted of peptidoglycan, the recovery of rod-shape through successive generations of growth was dependent upon the Rcs phosphorelay, in addition to PBP1B, LpoB and Lpp (34). It was proposed that unidentified members of the Rcs regulon could be responsible for the restoration of cell shape. Survival of L-form bacteria lacking a cell wall is also known to be dependent on RcsB, RcsC, RcsF and PBP1B (118).

Ongoing research is needed to uncover the precise role of the Rcs phosphorelay in round-cell mutants. However, we propose that in *Salmonella* Δ*pbpA* or Δ*rodA* mutants, in response to inherent cell wall defects RcsC activates genes required for survival in ‘stressful’ environments. Concurrently, virulence genes associated with intracellular survival are activated, whilst genes required for invasion in the nutrient-rich gut, are downregulated. It remains to be investigated whether the SPI-2 and Vi biosynthesis genes were upregulated in round-cell mutants, in an Rcs-dependent manner, although SPI-2 did not appear to be upregulated in *S.* Typhimurium Δ*mreC* cells (77). It is clear that the situation and regulatory controls involved are complex, and other regulators may contribute significantly to the observed phenotypes - as evidenced by the lack of clear SPI-1/flagella expression recovery in precise Δ*pbpA*/Δ*rodA* Δ*rcsC*/Δ*pagN* double-mutants.

The recovery of motile mutants harbouring transposon insertions within *pagN* further demonstrates the complexity of this system. Although only recovered once in the course of the screen, revertant mutants demonstrated a recovery in motility and SPI-1 protein expression. *pagN* is a SPI-6-encoded outer-membrane adhesin. The PagN invasin may compensate for a lack of SPI-1 activity, perhaps enabling *Salmonella* to escape from the host cell when SPI-1 is downregulated (119–122). *pagN* is co-regulated with SPI-2 by PhoP/PhoQ, and upregulated in the intracellular environment (98). Thus, it is potentially upregulated in Δ*pbpA* and Δ*rodA* mutants. To date, a potential role for PagN in virulence gene regulation has not been identified.

A crucial aspect of this study involved examining the virulence of the Δ*pbpA* and Δ*rodA* mutants *in vivo*, in the mouse model of typhoid fever infection. Surprisingly, both *S.* Typhimurium Δ*pbpA* and Δ*rodA* mutants inoculated intravenously were significantly attenuated *in vivo*, despite SPI-2 remaining actively expressed in these mutants *in vitro*. Significantly, this result was also contrary to a *S.* Typhimurium Δ*mreC* mutant which remained virulent *in vivo* (77).

In the complex *in vivo* environment multiple interconnected factors – many of which are not seen *in vitro* – may affect the survival and spread of these mutants. Specific *in vivo* environmental cues could lead to differential virulence gene regulation in the round-cell mutants, such that SPI-2 is downregulated. It is possible that alternative essential virulence factors are also inactivated as a result of the Δ*pbpA* or Δ*rodA* mutations, causing attenuation *in vivo*. In addition, inherent cell wall perturbations may simply render Δ*pbpA* and Δ*rodA* mutants unable to withstand the macrophage oxidative burst, to survive intracellularly. Unforeseen phenotypic variations between the *mrd* and *mre* mutants may explain the differences in the virulence of these strains *in vivo*. The present results may suggest that the cell wall defects resulting from Δ*pbpA* or Δ*rodA* knockouts are more severe.

Notably, the long time-course of the infection likely permitted the development of protective long-term immunity in the mice. This feature highlights the potential use of these mutant strains as vaccine candidates. Current licensed vaccines against typhoid fever have limited efficacy and are not available for very young children (122). Furthermore, the Vi subunit vaccine is ineffective against Vi-negative clinical strains of *S.* Typhi (123).

In conclusion, this study provides insights into the impact of the inactivation of *pbpA* and *rodA*, major members of the peptidoglycan-synthesis complex, on virulence in *Salmonella*. We have demonstrated that the inactivation of *pbpA* or *rodA* results in the specific downregulation of invasion-associated virulence genes. This is caused not directly by defects in the cell wall structure preventing virulence organelle function, but by the specific activity of the Rcs phosphorelay envelope stress response. In particular our results have highlighted the importance of RcsC in peptidoglycan synthesis mutants. Our findings in Δ*pbpA* and Δ*rodA* mutants reflect and validate those of a previous study in *S.* Typhimurium Δ*mreC* (77), further exhibiting the functional relationship between these genes.

This research is extremely relevant given that the bacterial envelope forms the immediate contact with the host, itself both comprising and supporting major virulence factors. Our future studies seek to further elucidate the regulatory mechanisms responsible for the observed virulence phenotypes in round-cell mutants, seeking to better understand both the roles of RcsC and PagN in virulence regulation, and the relationships between cell wall synthesis and virulence in Gram-negative pathogens. It also is recognised that post-transcriptional factors, such as the secondary messenger c-di-GMP, may play an important regulatory role – something which future work will explore. Further work is needed to investigate the activity of SPI-2 *in vivo* in Δ*pbpA* or Δ*rodA* mutants, and ultimately to assess the potential for *S.* Typhi or Typhimurium Δ*pbpA* or Δ*rodA* strains serving as much-needed vaccine candidates.

## Materials and Methods

### In silico analyses of the Salmonella mrd operon

Genomic DNA sequences of both *S.* Typhi and *S.* Typhimurium, within and immediately surrounding the putative *mrd* operon, were analysed and compared to published *E. coli* datasets. The following *in silico* tools were used for these analyses: Basic Local Alignment Search Tool (BLASTn/megablast - http://blast.ncbi.nlm.nih.gov/Blast.cgi) (58), RegulonDB (http://regulondb.ccg.unam.mx/) (124), BDGP Fruitfly (www.fruitfly.org) (55) and Softberry BPROM (www.softberry.com) (56) (promoter prediction tools), and TranstermHP (http://transterm.cbcb.umd.edu/) (57) (transcriptional terminator algorithm).

### Strains and culture conditions

All strains (Table 1) were routinely cultured in Luria-Bertani (LB) broth containing relevant antibiotics (ampicillin: 100 µg ml^−1^, kanamycin: 50 µg ml^−1^, chloramphenicol: 25 µg ml^−1^, tetracycline: 50 µg ml^−1^), at 37°C and 200rpm. *S.* Typhi strain BRD948 cultures were supplemented with aromatic amino acids and tyrosine, as described (125). Δ*pbpA* and Δ*rodA* strains were grown in media containing 20mM MgCl_2_.

**Table 1 –.**
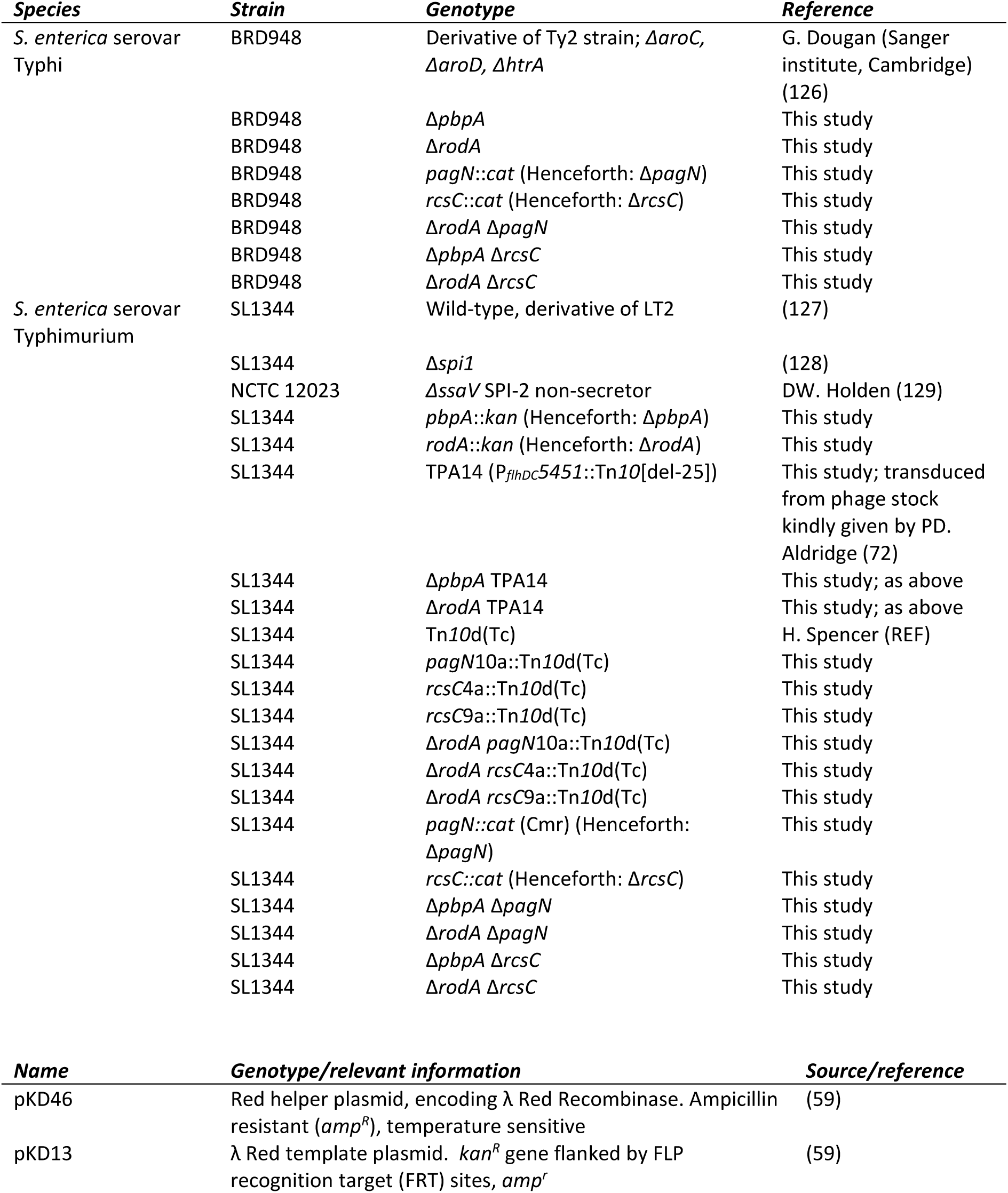
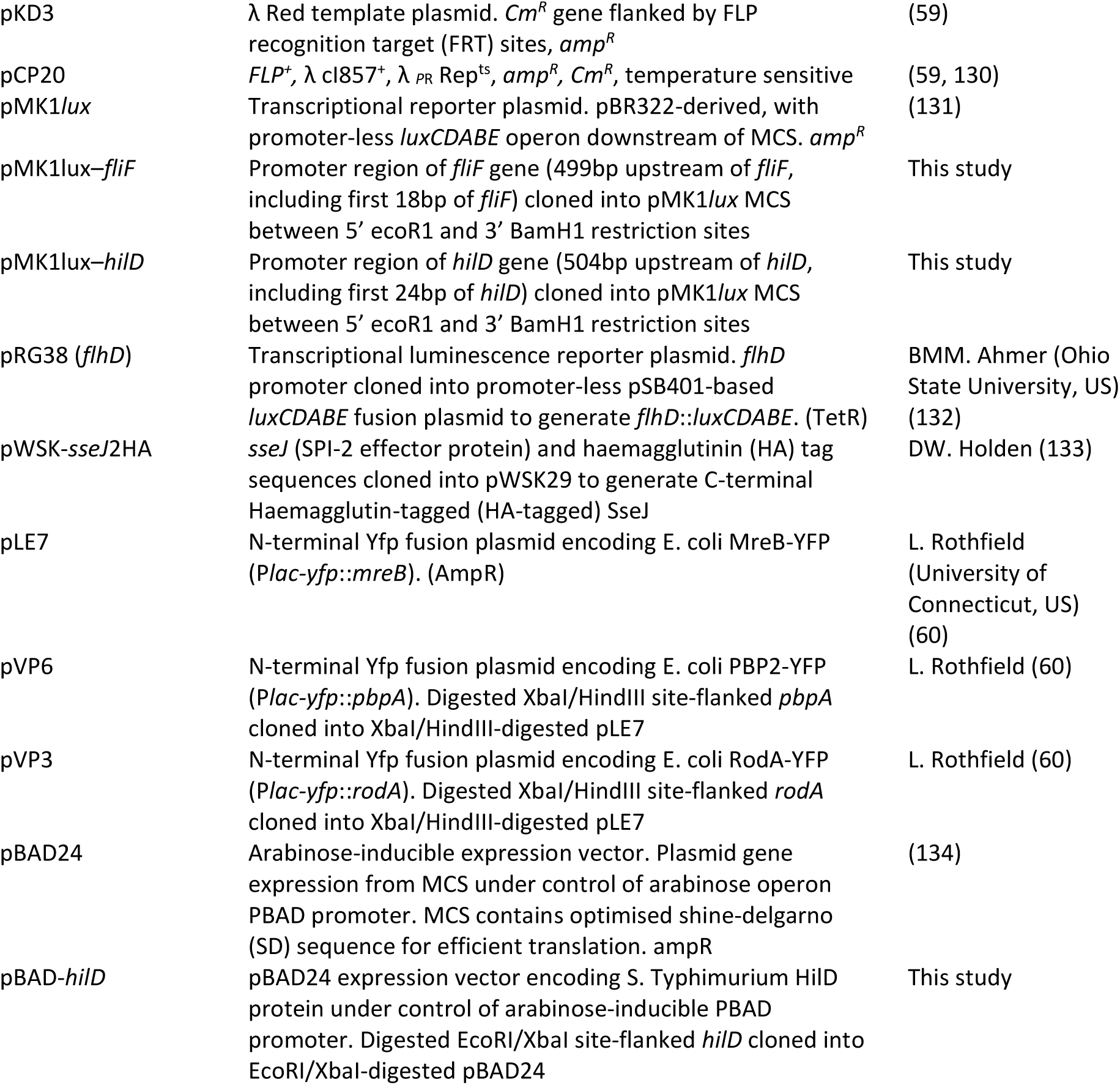
Strains and Plasmids.

5ml cultures were grown overnight before subculturing 1:100 into 25-50ml fresh LB (250ml flasks). Cultures were harvested at mid-log phase (OD600_nm_ ~1.2) for the generation of electrocompetent cells and studies in SPI-1-inducing conditions. For SPI-2 expression studies, overnight cultures were subcultured 1:100 into SPI-2-inducing minimal media, as described (77), without additional MgCl_2_. Cultures were grown for 16 hours at 37°C and 200rpm before harvesting.

**Table 2 –.**
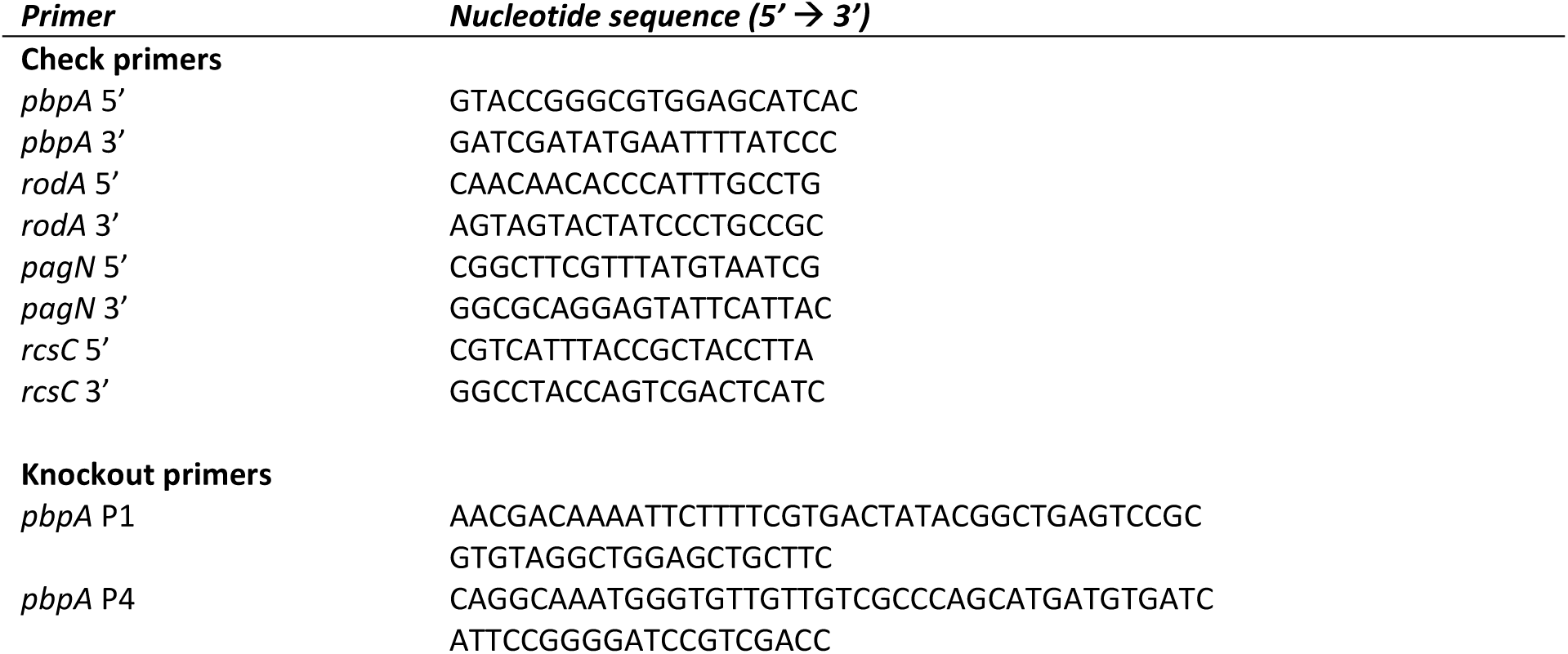
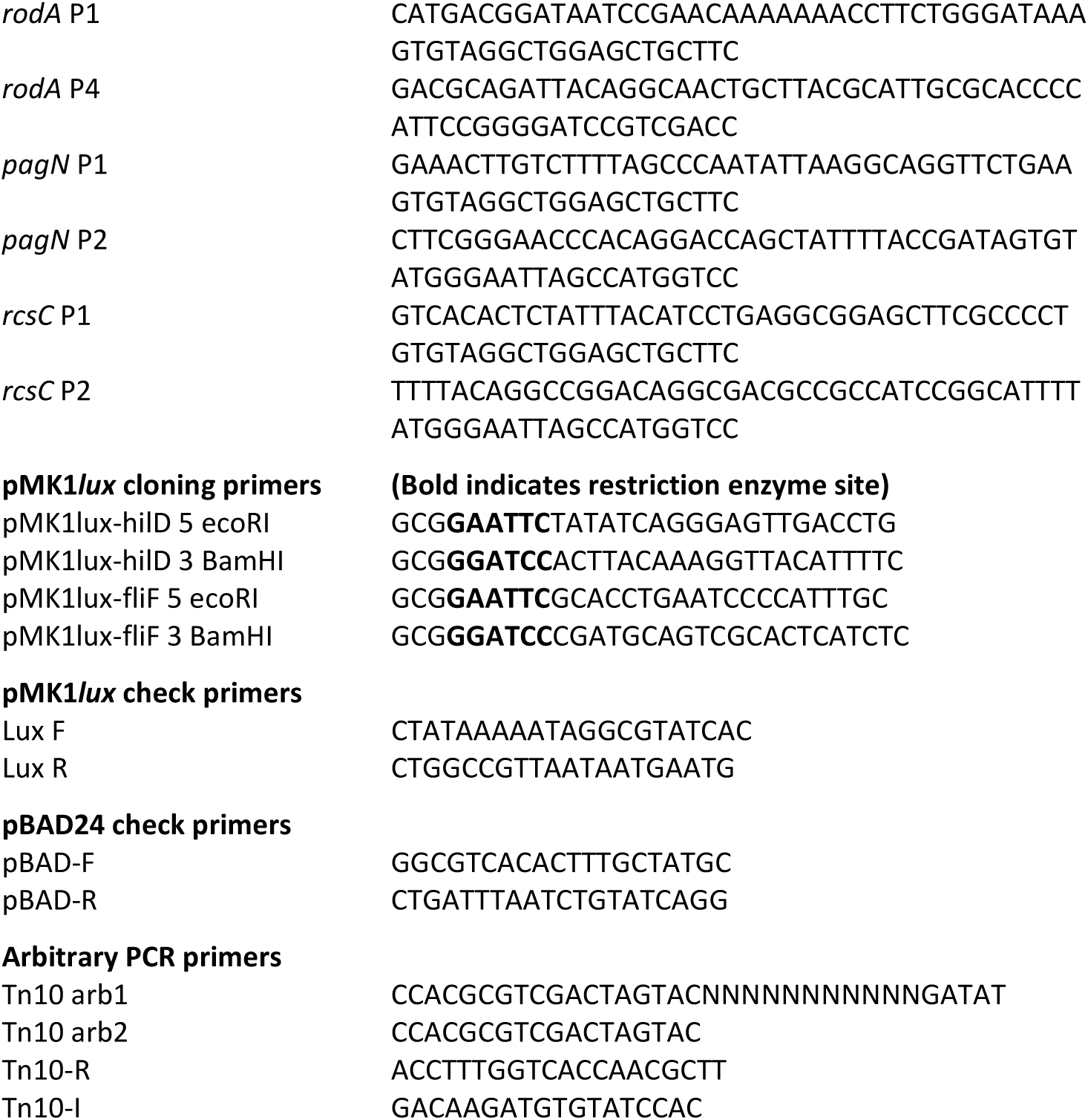
PCR Primers.

### Generation and complementation of chromosomal gene disruptions in *Salmonella*

Chromosomal deletion mutants of *pbpA*, *rodA*, *rcsC* and *pagN* were generated according to the Lambda-Red method, as previously described (59). All washes during the generation of electrocompetent cells were carried out in ice-cold sterile water containing MgCl_2_ (20mM). Complete knockout mutants of *pbpA* and *rodA* were generated in *S.* Typhi BRD948, leaving a short 82bp scar sequence (a single FRT site flanked by short 5’ and 3’ sections of the wild-type gene). We were unable to remove the kanamycin resistance cassette from Δ*pbpA* and Δ*rodA* mutants of *S.* Typhimurium SL1344. Chloramphenicol-resistant non-polar BRD948 and SL1344 Δ*rcsC* and Δ*pagN* mutants were constructed using the pKD3 template plasmid. Successful generation of each mutant was checked by PCR and DNA sequencing.

### Motility assays

1-2 μl fresh overnight *Salmonella* cultures were inoculated into semi-solid agar (10 g l^-1^ tryptone, 5 g l^-1^ NaCl and 3 g l^-1^ agar). Plates were incubated upright at 37°C for 5-6 hours, after which time the extent of culture spread (diameter) from the site of inoculation was measured, as compared to both wild-type and non-motile controls.

### Preparation of protein fractions for analysis of protein expression

To prepare whole-cell protein fractions, bacteria were harvested from liquid culture by centrifugation and resuspended in an appropriate volume PBS, according to the culture optical density (OD600_nm_). Bacterial secreted proteins were precipitated from filter-sterilised culture supernatants (pore size – 0.22μm) by adding 10% trichloroacetic acid and incubating for 16 hours at 4°C. Secreted proteins were then concentrated by centrifugation (12000 x g for 30 minutes) and resuspension in 300 μl 2% SDS and 1.2 ml acetone. Samples were stored at −80°C for 16 hours, pelleted and finally resuspended in ≥25 µl PBS.

SDS loading buffer (containing mercaptoethanol) was added to protein samples, which were then boiled for 10 minutes before being loaded onto 12% acrylamide gels for separation by SDS-PAGE.

For Western Blotting, proteins were transferred onto Protran nitrocellulose membranes (Schleicher & Schuell) using a wet transfer apparatus (Biorad). Membranes were probed using polyclonal αSipA, αSipC, αSipD or αSopE (SPI-1), or αSseB (SPI-2) antibodies (kindly provided by Vassilis Koronakis, Cambridge, UK), coupled with a goat anti-mouse horseradish peroxidase-labelled secondary antibody (Dako Cytomation). To test for SPI-2 functionality membranes were probed first with monoclonal HA.11 IgG antibody (Covance Inc., New Jersey). Membranes were developed using 4-chloro-1-naphthol. These assays were repeated at least three times using various different effector protein antibodies (SPI-1).

### Microscopy

Phase-contrast and fluorescence microscopy images were viewed and captured under a 100x/NA 1.4 oil immersion objective, using a Nikon Ti-E microscope coupled to an Andor iXon^EM^+ 885 EMCCD camera. Both NIS-ELEMENTS (Nikon) and ImageJ software were used to process microscopy images.

### Visualisation of SPI-1 and SPI-2 T3S needles by immunofluorescence microscopy

*Salmonella* cultures grown in SPI-1 or SPI-2-inducing conditions were harvested and fixed in 4% Paraformaldehyde (in PBS) for 30-60 minutes. Cells were then washed for 15 minutes with three successive PBS washes, before being incubated for 1 hour in primary αSipD (SPI-1) or αSseB (SPI-2) antibody diluted 1:1000 in PBS. After washing as before, cells were probed with Alexa Fluor 568-conjugated goat anti-rat antibody or 488-conjugated goat anti-rabbit antibody (Invitrogen-Molecular Probes, Paisley, U.K.), diluted 1:200 or 1:1000 in PBS, respectively. Samples were incubated in the dark for 2 hours and washed as before. Cells were finally resuspended in 100-300 μl 10% glycerol and mounted onto thin 1% agarose beds. Phase contrast and immunofluorescence microscopy images were taken of the same field and then merged.

### Vi slide agglutination assays

In order to assess Vi capsule production in *S.* Typhi strains, 5-10 μl fresh overnight BRD498 cultures (plus WT SL1344 as a negative control) were individually dropped onto clean microscope slides. An equal volume of Vi-agglutinating serum (Murex Biotech Ltd, Dartford, UK) and sterile PBS were then added and gently mixed. Slides were incubated at 22°C for several minutes and then observed for signs of agglutination.

### Transcriptional reporter fusion assays

Promoter regions of the *S.* Typhi *fliF* and *hilD* genes were individually cloned into the promoter-less *lux* operon-encoding transcriptional reporter plasmid, pMK1*lux* (131). Successful cloning was verified by sequencing. pMK1*lux* transcriptional reporter fusions were transformed into WT, Δ*pbpA* and Δ*rodA* strains (BRD948 and SL1344). Overnight cultures of the resulting strains were diluted 1:1000 into fresh LB (with appropriate supplements) and grown for 16 hours in microtitre plates (200 μl volume) at 37°C, with shaking, in a Tecan Infinity200 luminometer. Relative luminescence emitted from the cultures (luminescence/OD600_nm_) was recorded at 15 minute intervals during the growth cycle, as a measure of *lux* operon expression. Assays were conducted in triplicate with at least 3 independent repeats.

### Tn*10*d(Tc) transposon mutagenesis screen

Tn*10*d(Tc) is a 2.64 kbp transposon-defective mini transposon derived from Tn*10*, which requires the exogenous expression of a plasmid-borne transposase (135, 136). A P22 phage lysate library pooled from 12000 SL1344 Tn*10*d(Tc) transductant colonies was kindly provided for use in the mutagenesis screen (Hannah Spencer, Newcastle). Overnight cultures of SL1344 Δ*rodA* pMK1*lux-fliF* were transduced with the Tn*10*d(Tc) phage lysate, as previously described (137). After transduction cells were recovered in warm LB before being spread onto LB agar containing tetracycline and incubated overnight at 37°C. To identify transductants with recovered *fliF* expression, tetracycline-resistant transductant colonies, along with appropriate positive and negative controls, were patched in duplicate onto fresh LB agar containing ampicillin. After growth overnight at 37°C, one plate was blotted onto damp filter paper, covered in transparent film and exposed to light-sensitive film in the dark for 20-30 seconds. Luminescent colonies – evident as dark spots on the developed film – were re-patched and assessed again for recovered luminescence. Arbitrary PCR and sequencing was then performed on positive revertants, as described previously (137–139), in order to map the genomic location of the Tn*10*d(Tc) insertions. Tn*10*d(Tc) lesions from the positive revertants were each transduced back into the parent SL1344 WT/Δ*rodA* strain using the P22 phage, prior to further phenotypic analysis.

### Ethics statement

*in vivo* experiments were covered by a Project License, granted by the Home Office under the Animal (Scientific Procedures) Act 1986. This license was approved locally by the University of Cambridge Ethical Review Committee.

### *in vivo* inoculation of mice & growth curves

*in vivo* virulence assays were conducted in C57BL/6 mice as previously described (77). SL1344 WT and Δ*rodA* strains were inoculated intravenously into mice in order to bypass the SPI-1-dependent stages of infection. The liver and spleen bacterial loads were subsequently measured at 6 hours, 1 day, then at 3, 6, 14 and 28 days post-infection (colony-forming units (CFU) per organ). Average results from 4 mice were recorded. Where possible, all assays were conducted out in triplicate, with three independent repeats.

## Acknowledgements

We thank Laurence Rothfield and Brian Ahmer for generously providing the vectors pVP6, pVP3, and pRG38. ACD, BCG and JCC were supported by MRC DTP and BBSRC DTP Studentships awarded to CMAK.

